# Backbone Brackets and Arginine Tweezers delineate Class I and Class II aminoacyl tRNA synthetases

**DOI:** 10.1101/198846

**Authors:** Florian Kaiser, Sebastian Bittrich, Sebastian Salentin, Christoph Leberecht, V. Joachim Haupt, Sarah Krautwurst, Michael Schroeder, Dirk Labudde

## Abstract

The origin of the machinery that realizes protein biosynthesis in all organisms is still unclear. One key component of this machinery are aminoacyl tRNA synthetases (aaRS), which ligate tRNAs to amino acids while consuming ATP. Sequence analyses revealed that these enzymes can be divided into two complementary classes. Both classes differ significantly on a sequence and structural level, feature different reaction mechanisms, and occur in diverse oligomerization states. The one unifying aspect of both classes is their function of binding ATP. We identified Backbone Brackets and Arginine Tweezers as most compact ATP binding motifs characteristic for each Class. Geometric analysis shows a structural rearrangement of the Backbone Brackets upon ATP binding, indicating a general mechanism of all Class I structures. Regarding the origin of aaRS, the Rodin-Ohno hypothesis states that the peculiar nature of the two aaRS classes is the result of their primordial forms, called Protozymes, being encoded on opposite strands of the same gene. Backbone Brackets and Arginine Tweezers were traced back to the proposed Protozymes and their more efficient successors, the Urzymes. Both structural motifs can be observed as pairs of residues in contemporary structures and it seems that the time of their addition, indicated by their placement in the ancient aaRS, coincides with the evolutionary trace of Proto- and Urzymes.

**Author summary:** Aminoacyl tRNA synthetases (aaRS) are primordial enzymes essential for interpretation and transfer of genetic information. Understanding the origin of the peculiarities observed with aaRS can explain what constituted the earliest life forms and how the genetic code was established. The increasing amount of experimentally determined three-dimensional structures of aaRS opens up new avenues for high-throughput analyses of molecular mechanisms. In this study, we present an exhaustive structural analysis of ATP binding motifs. We unveil an oppositional implementation of enzyme substrate binding in each aaRS Class. While Class I binds via interactions mediated by backbone hydrogen bonds, Class II uses a pair of arginine residues to establish salt bridges to its ATP ligand. We show how nature realized the binding of the same ligand species with completely different mechanisms. In addition, we demonstrate that sequence or even structure analysis for conserved residues may miss important functional aspects which can only be revealed by ligand interaction studies. Additionally, the placement of those key residues in the structure supports a popular hypothesis, which states that prototypic aaRS were once coded on complementary strands of the same gene.

## Introduction

The synthesis of proteins is fundamental to all organisms. It requires a complex molecular machinery of more than 100 entities to ensure efficiency and fidelity [1–3]. The ribosome pairs an mRNA codon with its corresponding anticodon of a tRNA molecule that delivers the cognate amino acid. Aminoacyl tRNA synthetases (aaRS) ligate amino acids to their corresponding tRNA [4], which is why they are key players in the transfer of genetic information. The mere existence of proteins and nucleic acids is a chicken or the egg dilemma. The sequential succession of amino acids in each protein is encoded by nucleic acid blueprints. In turn, these proteins are indispensable to replicate and translate nucleic acids. It is debated how this reflexive system came to be [5] and which polymer type constituted the earliest living systems. The RNA world hypothesis assumes nucleic acids were the sole basis of primordial life. RNA molecules can store and interpret genetic information, while also allowing for catalytic activity. In succession, proteins emerged to implement more elaborate, specific, and efficient catalytic activity [6]. However, the limited catalytic repertoire of RNA molecules [7] raises concerns that such a primordial world was based on a single polymer type. The peptide-RNA world hypothesis assumes that life and genetic information originated from a system in which RNA and peptides coexisted and complemented each other from the very beginning [7–10]. It is argued that only this interleaving of the two types of macromolecules can account for the speed with which the genetic code developed [9, 11–13]. Both hypotheses were reviewed recently [11, 14]. Either way, aaRS are the entities which most prominently reflect that early episode of life.

The unique interface between gene and gene products is shaped by aaRS as they attach the amino acid to the corresponding tRNA molecule [4, 11]. Three main theories have been proposed to explain the emergence of the self-encoding translational machinery, namely: coevolution [15], ambiguity reduction [16, 17], and stereochemical forces [18]. The interaction between amino acid and nucleic acid lies at the basis of each theory and is linked to the emergence of aaRS [7, 19]. There is strong evidence for two archaic proto-enzymes as the origin of all aaRS, which were among the earliest proteins that enabled the development of life [8, 20–22]. Since then, these predecessors have evolved divergently into Class I and Class II (Figure 1), where each is responsible for a distinct set of amino acids [23–25]. The physicochemical properties of amino acids are distributed evenly between both classes, even though amino acids handled by Class I were shown to be slightly bigger [26]. This suggest a concurrent emergence of both classes and that archaic aaRS substrates have differed sufficiently to require two specialized kinds of aaRS [11]. Both classes are, on several levels, as distinct as possible from each other [11].

**Fig 1.**
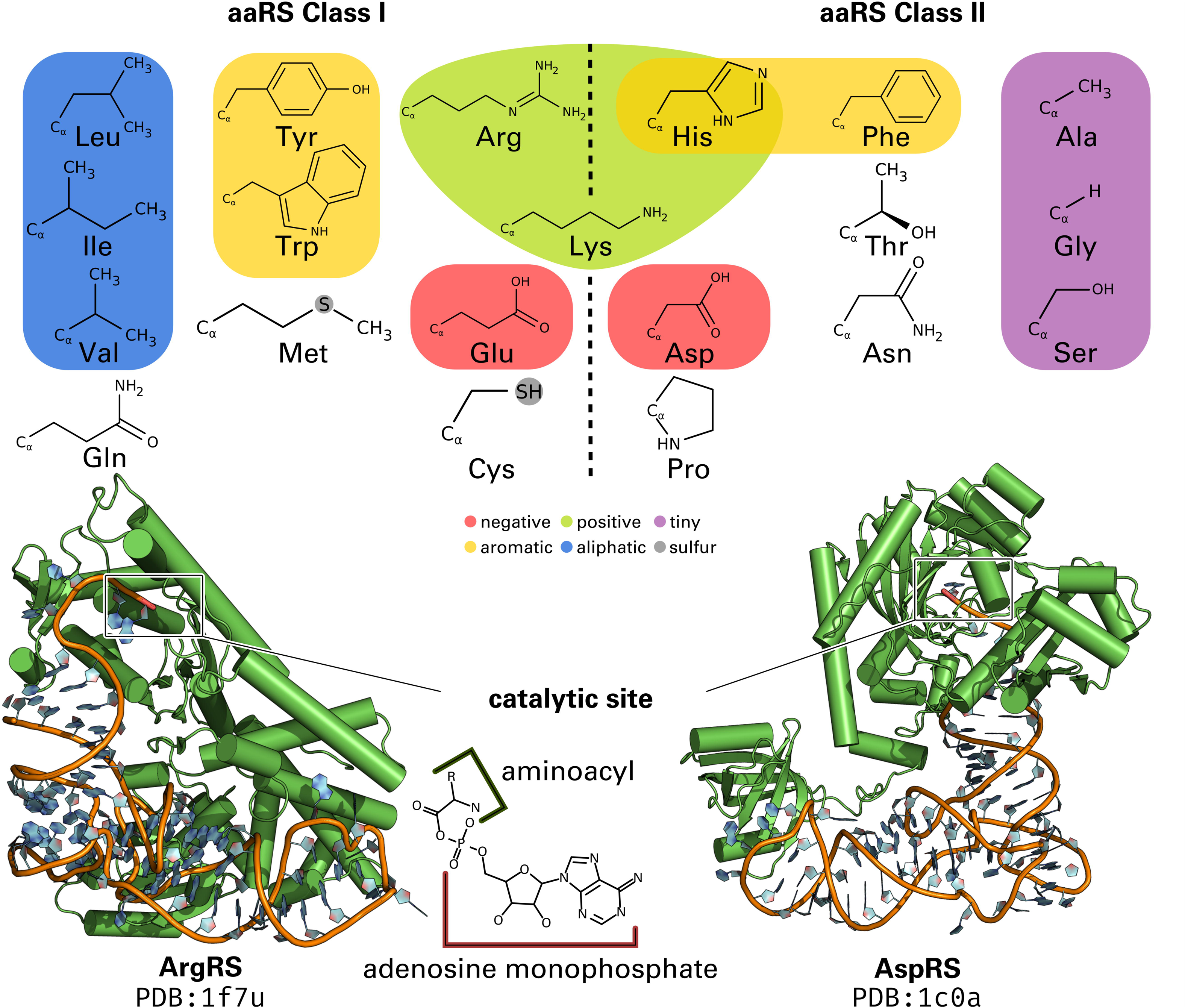
The two aaRS classes and amino acids they ligate to the cognate tRNA. Based on the physicochemical properties of the amino acids (colored according to [40]) no distinction can be made between the two classes. However, statistically significant differences based on amino acid side chain size [26] and binding site size [41, 42] are evident. Lysine is mostly processed by Class II aaRS, but in all archaic organisms a Class I aaRS is responsible for lysine [43]. Prior to tRNA ligation, the amino acid ligand is converted to its activated form: aminoacyl adenylate.

Every aaRS recognizes an amino acid and prevents misacylation of tRNAs by maximizing ligand specificity. The discrimination mechanisms between similar amino acids are well-studied [4, 27–29]. During the enzymatic reaction the designated amino acid is activated, forming an aminoacyl adenylate, before it is linked to the cognate tRNA [30, 31]. For example, the fusion of aspartic acid and its corresponding tRNA^Asp^ by the aspartyl-tRNA synthetase (AspRS) follows the two-step reaction:

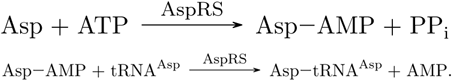

Today most organisms feature 20 concrete realizations, each handling one specific amino acid [32, 33] – throughout the paper they are referred to as aaRS Types.

The modular architecture of aaRS has evolved well-orchestrated and was optimized for its specific requirements [24, 34]. Frequent domain inserts [9, 11] can render the evolutionary origin hard to track [35]. In principle, all aaRS have to conserve three functions: the correct recognition of the tRNA identity and amino acid as well as the ligation of both. Commonly, the anticodon binding domain ensures tRNA integrity by recognizing particular features of the anticodon [36, 37]. The identification and transfer of amino acids is then mediated by the catalytic domain, which differs in topology between the two classes (Fig 1). To minimize errors in protein biosynthesis, pre- and post-transfer editing mechanisms are conducted by approximately half of the aaRS Types [27, 38, 39].

Sequences of aaRS proteins are highly diverse and result from fusion, duplication, recombination, and horizontal gene transfer [44, 45]. However, two sets of Class-specific and mutually exclusive sequence motifs have been identified, which are responsible for interactions with adenosine phosphate as well as catalysis [4, 23, 46]. Class I features the conserved HIGH and KMSKS motifs [4, 23]. The functional key motifs in Class II are referred to as Motif “1”, Motif “2”, and Motif “3” [4]. Both HIGH and KMSKS stabilize the transition state, whereby the latter constitutes a mobile loop in the folded structure [4]. The binding of ATP and the transition state of the reaction of individual Class I proteins have been demonstrated to be stabilized by a structural rearrangement [8, 47–54], which stores energy in a constrained conformation of the KMSKS motif [55]. The Class II motifs are less conserved [35] and more variable in their relative arrangement [23]. Motif “1” mediates the dimerization of protein structures, commonly found in Class II aaRS [4, 56]. Motif “2” and “3” are essential for the reaction mechanism and feature two highly conserved arginine residues [23, 57, 58].

The catalytic domain of Class I adapts a Rossmann fold [30], whereas Class II possesses a unique fold [45, 59, 60]. To assert the global structural similarity, two major structural alignments were calculated for Class I and Class II, respectively, that revealed high structural similarity within each Class with average sequence identity below 10% [61]. On a functional level, both aaRS classes exhibit distinct ATP binding site architectures and reaction mechanisms. Class I aaRS proteins attach the amino acid to the 2’OH-group of the tRNA’s 3’-terminal adenosine, whereas Class II proteins use the 3’OH-group as the attachment location [62].

## Rodin-Ohno hypothesis

In 1995, Rodin and Ohno proposed an elegant explanation for the peculiarities that are observed in contemporary aaRS: both classes were originally encoded on complementary strands of the same nucleotide fragment [8] (Fig 2). The Rodin-Ohno hypothesis is supported by an experimental deconstruction of aaRS sequences [9, 11]. In these studies, parts of contemporary aaRS proteins were removed and the catalytic strength of the resulting transcripts was assessed. One representative sequence of each Class was reduced to a peptide of only 46 amino acids. The coding nucleotide sequences of these 46-residue peptide were paired complementarily. These so called “Protozymes” were investigated regarding their structural and catalytic properties; they form molten globules [9, 11] and – despite the lack of ordered tertiary structure – they are still capable of rate enhancements by orders of magnitude [9, 11]. It is essential that the efficiency of different enzyme families across the proteome increases at comparable rates [9, 11]. The phenomenon of anti-parallel coupling of two genes was also postulated for other families of proteins [63, 64] and seems to be a phenomenon that affects the whole genome [65, 66]. One contradicting theory is the coevolutionary theory of the genetic code [15]. This theory suggests two main groups of amino acids based on the connectedness of their biochemical pathways and that amino acid biosynthesis was the dominant factor that shaped the genetic code [19]. Other authors suggested that both classes evolved from unrelated ancestors and are of independent origin [23].

**Fig 2.**
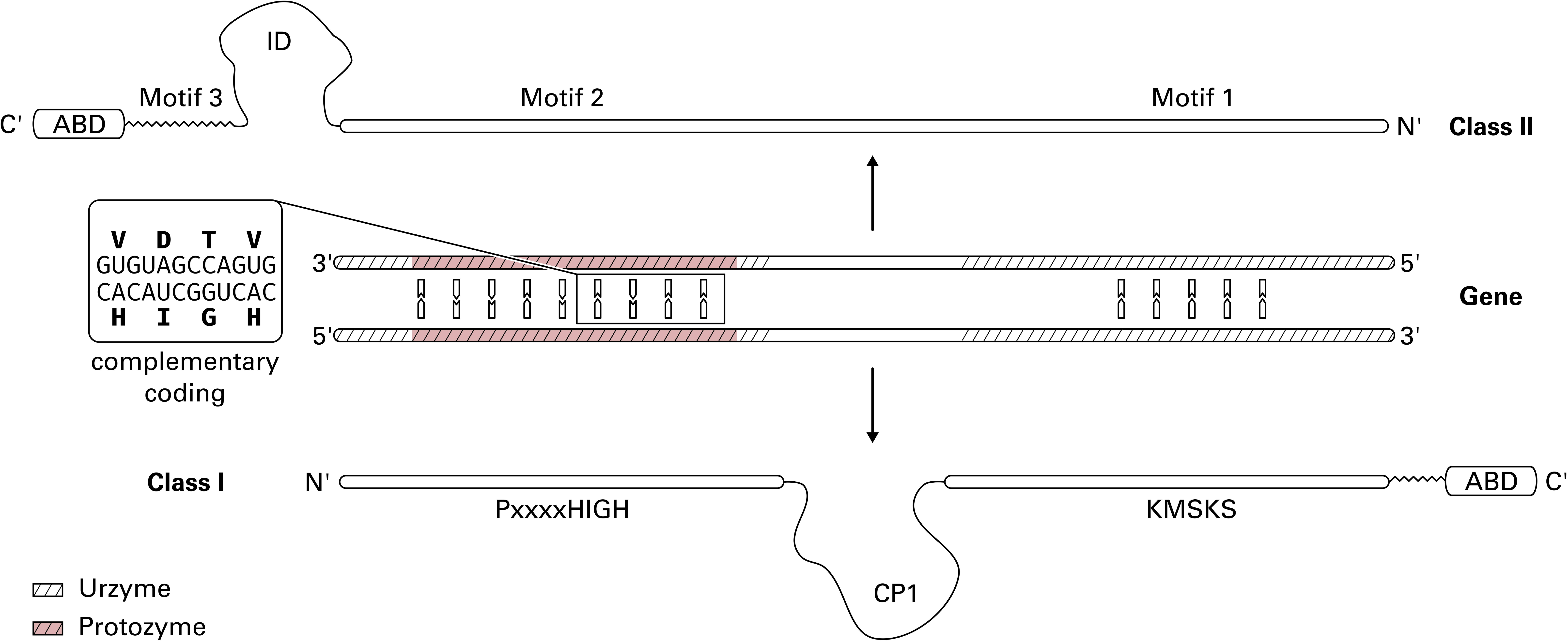
The Rodin-Ohno hypothesis states that both aaRS classes descended from the opposite strands of a single gene. The signature motifs of each class were fully complementary on this gene. Both Protozymes originated from the complementary “HIGH-Motif 2” region (shaded in red). Contemporary aaRS feature insertion domains (ID) and Connecting Peptides (CP1) as well as the addition of the anticodon binding domain (ABD). Figure adapted from [9, 67].

The Rodin-Ohno hypothesis can explain why ATP and tRNA binding sites of both classes seem to be mirror images of each other [68] as well as the fact that both classes share virtually no similarities [4, 11, 45] beside their actual function [8, 9, 11]. All of the contemporary aaRS Types are connected by the requirement to bind ATP. This basal unifying characteristic was found to involve hydrogen bonds in the Class I Protozymes [9].

Remarkably, the restrictions inherent with a complementary coding may explain why the middle base of a codon is the most distinctive base for the corresponding amino acid nowadays [22]. Other studies showed how slight differences in the substrate can result in a stable separation of aaRS into two classes [7, 10]. Potentially, the two Protozymes diverged into ten aaRS Types each (Fig 1) and simultaneously increased fidelity and incorporated additional domains when necessary [8, 9, 20–22]. Most of aaRS evolution took place before the “Darwinian threshold” [61]. Only a small number of amino acids, such as tryptophan, were gradually incorporated into the genetic code after the last universal common ancestor and inefficient proteins evolved over time [19]. While similar amino acids were once processed by the same aaRS, specificity may have required additional aaRS Types to cope with increasing complexity. It is still possible to observe such generic aaRS in some organisms [69, 70].

## Motivation

A systematic delineation of aaRS active site residues is expedient [11]. The most conserved part of the aaRS reaction mechanism is the amino acid activation with ATP, since it represents the principal kinetic barrier for the creation of peptides in a pre-biotic context [67]. This fundamental mechanism is shared by all Class I and Class II aaRS enzymes, irrespective of their Type or the organism of origin. Furthermore, the catalytic domain has been predicted to constitute the ancestral aaRS precursors [9, 11, 64, 71].

The residues of the catalytic domain involved in amino acid binding were molded to meet specific requirements of the individual properties of each amino acid during evolution. In contrary, the ATP binding part includes the most conserved parts of the structure. To achieve a systematic delineation of available protein structures, this study focuses on the most common element: the binding of the ATP substrate. Individual aaRS and their mechanism to discriminate similar amino acids have been extensively studied on the structural level [4, 27–29]. However, a comprehensive and comparative study of structural features in aaRS proteins is missing. There are no structural motifs known that capture the profound differences of the ligand recognition mechanism.

To unveil general adenosine phosphate binding properties of each aaRS Class, we have investigated the corresponding binding pockets of 972 aaRS protein molecules for each aaRS Type across all kingdoms of life. In total, 448 protein chains for Class I and 524 chains for Class II, available from the Protein Data Bank (PDB) [72], were analyzed. Previous studies have focused on comparing subsets of structures for each Class but to our knowledge no conclusive study was conducted that includes structures for every aaRS Type for both classes.

The results of this study outline the dichotomy between the two classes (Fig 3) on a functional level. A conserved pair of arginine residues is grasping the adenosine phosphate part of the ligand in nearly all Class II structures. Class I features no comparable structural pattern for adenosine phosphate binding, but interaction analysis divulged two highly conserved backbone hydrogen bonds, which seem to realize the same function without the need for conserved amino acid side chains. Due to their geometrical characteristics, we refer to the Class I and Class II motifs as Backbone Brackets and Arginine Tweezers, respectively. The Backbone Brackets motif demonstrates the limitations of sequence analysis and was, to our knowledge, never identified as a highly conserved interaction pattern prior to this study. Additionally, a novel geometrical characterization of these structural motifs demonstrates that significant structural rearrangements can be observed for all Class I structures upon ligand binding. The highly sensitive geometric characterization of side chain angles and alpha carbon distances is able to detect subtle differences in ligand binding and is potentially suitable to be applied on other conserved structural patterns as well.

**Fig 3.**
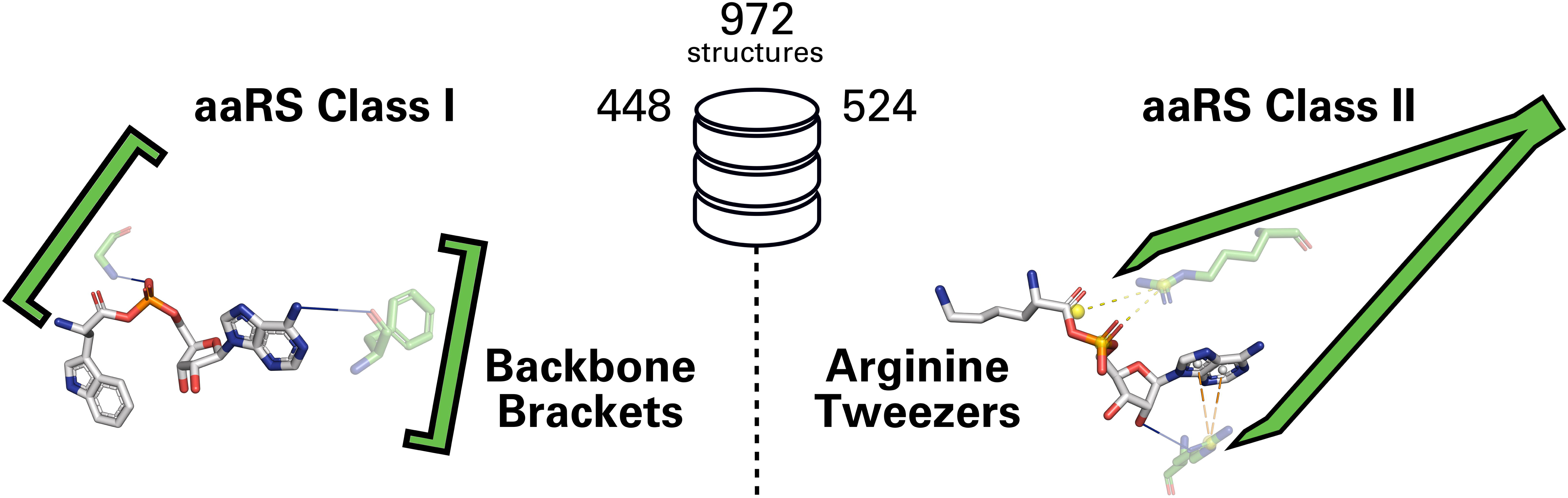
Backbone Brackets and Arginine Tweezers. Based on the analysis of 972 protein 3D structures (448 protein chains for Class I and 524 chains for Class II), Backbone Brackets and Arginine Tweezers were identified as structural motifs distinctive for their respective aaRS Class.

Both structural motifs can be traced back to the Protozyme and Urzyme regions postulated in the studies based on the Rodin-Ohno hypothesis [8, 9, 11]. The analysis of codons in the corresponding regions accentuates existing insights and allows for an additional look behind the curtain of evolution.

## Results

This study presents a dataset of aaRS structures annotated with ligand information, which serves as a stepping stone to understand common and characteristic ligand interaction properties. It is composed of 972 individual chains containing 448 (524) Class I (Class II) catalytic aaRS domains and covers at least one ligand-bound structure for each aaRS type. The dataset is provided in S1 File and S2 File. The Class I chains originate from 256 biological assemblies and comprise 151 bacterial, 84 eukaryotic (including four mitochondrial structures), 20 archaea, and one viral structure. The Class II chain set corresponds to 267 biological assemblies where 102 are of bacterial origin, 104 from eukaryotes (including 15 mitochondrial structures), and 61 from archaea. For a detailed organism overview see S9 Fig. The sequence identity is below 33% (29%) for 95% of all Class I (Class II) structures, while pairwise structure similarity is high with a TM score [73] over 0.8 for 95% of the structures (S8 Fig). The high sequential diversity probably stems from the variety of covered organisms and domain insertions. In contrast, the low structural diversity can be seen as a result of conserved function and the shared topology of the catalytic domain within each aaRS Class.

Sequence positions of all structures in the dataset were unified using a multiple sequence alignment (MSA) generated with the T-Coffee expresso pipeline [74] (Section Mapping of binding sites, S5 File and S6 File). This type of MSA is backed by the additional structural alignment of protein structures. Hence, the structurally conserved catalytic core region is preferred during alignment, since insertion domains and structurally diverse attachments do not align structurally across the whole dataset. The MSA allows the investigation of a plethora of structures independently of the concrete aaRS Type. This investigation is aided by a renumeration that effectively provides a means to compare sequentially divergent, structurally similar proteins. All further referenced positions are given in accordance to this MSA. In figures where depictions of structures are shown, the original sequence positions of residues are listed. To infer original sequence positions from given renumbered sequence positions, tables S13 File and S14 File are provided. These tables contain the corresponding original sequence positions for each position of the MSA and for each structure in the dataset.

### Backbone Brackets and Arginine Tweezers

In order to investigate the contacts between aaRS residues and their ligands, noncovalent protein-ligand interactions were annotated. This revealed two highly consistent interaction patterns between catalytic site residues and the adenosine phosphate part of the ligand: conserved backbone hydrogen bonds in Class I as well as two arginines with conserved salt bridges and side chain orientations in Class II.

Strikingly, the residues mediating the backbone interactions were mapped in 441 of 448 (98%) Class I renumbered structures at the two positions 274 and 1361. Closer investigation on the structural level revealed geometrically highly-conserved hydrogen bonds between the peptide bond’s nitrogen or oxygen atom and the adenosine phosphate part of the ligand (Fig 4A). These two residues mimic a bracket-like geometry (Fig 4B), enclosing the adenosine phosphate, and were thus termed Backbone Brackets. The interacting amino acids are not limited to specific residues as their side chains do not form any ligand contacts. Hence, position 274 of the Class I motif is not apparent on sequence level while position 1361 exhibits preference for hydrophobic amino acids, e.g. leucine, valine, or isoleucine (Fig 4C). Examples for the Backbone Brackets motif are residues 153 (corresponding to renumbered residue 274) and 405 (corresponding to renumbered residue 1361) in Class I ArgRS structure PDB:1f7u chain A.

**Fig 4.**
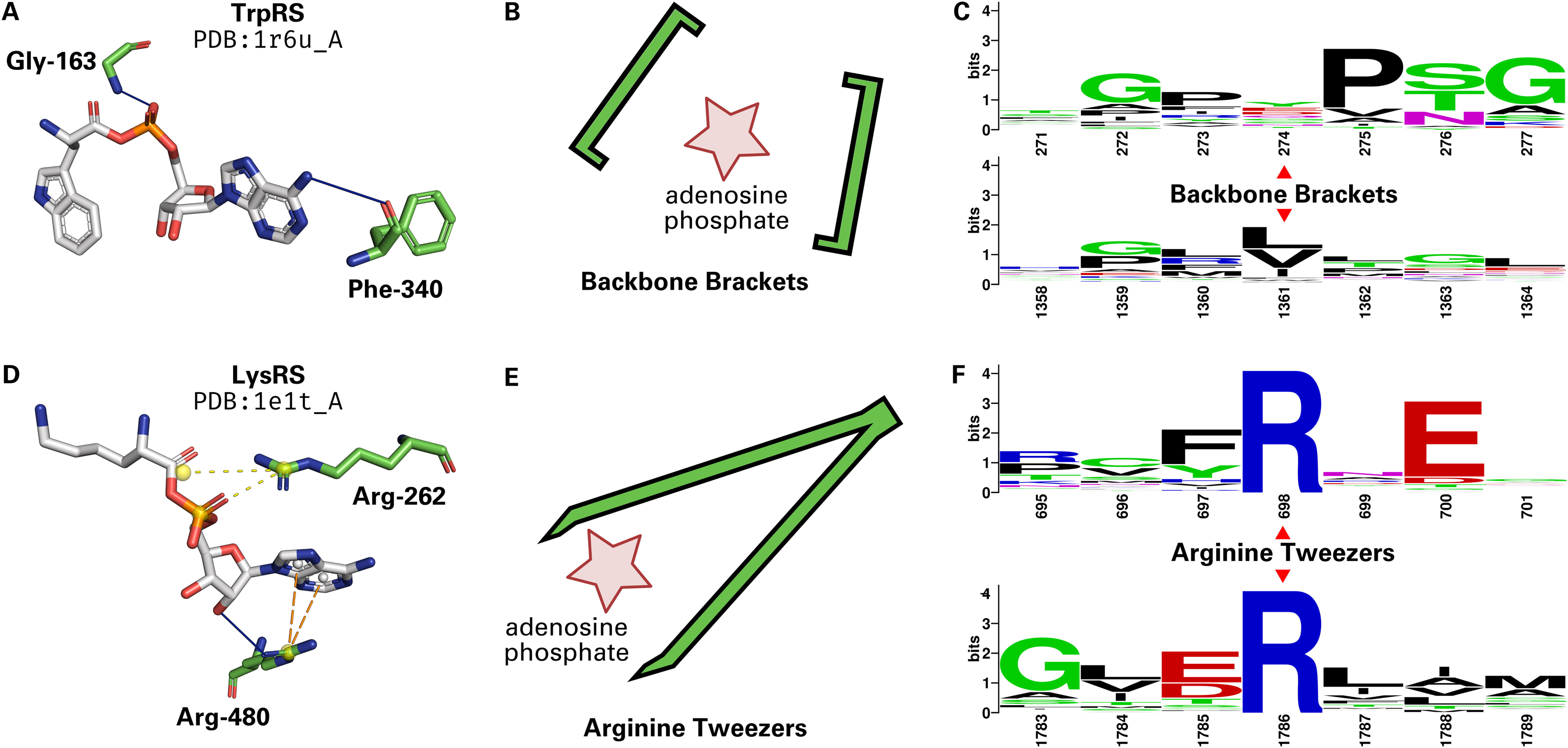
Comparison of Backbone Brackets and Arginine Tweezers. (A) Structural representation of the Backbone Brackets motif interacting with Tryptophanyl-5’AMP ligand in TrpRS (PDB:1r6u chain A). The ligand interaction is mediated by backbone hydrogen bonds (solid blue lines). Residue numbers are given in accordance to the structure of origin. (B) The geometry of the Backbone Brackets motif resembles brackets encircling the ligand. (C) WebLogo [75] representation of the sequence of Backbone Brackets residues (274 and 1361) and three surrounding sequence positions. Residue numbers are given in accordance to the MSA. (D) Structural representation of the Arginine Tweezers motif in interaction with Lysyl-5’AMP ligand in LysRS (PDB:1e1t chain A). Salt bridges (yellow dashed lines) as well as π-cation interactions are established. Residue numbers are given in accordance to the structure of origin. (E) The Arginine Tweezers geometry mimics a pair of tweezers grasping the ligand. (F) Sequence of Arginine Tweezers residues (698 and 1786) and surrounding sequence positions. The Backbone Brackets show nearly no conservation on sequence level since backbone interactions can be established by all amino acids, while the Arginine Tweezers rely on salt bridge interactions, always mediated by two arginines. Residue numbers are given in accordance to the MSA.

In contrast, Class II aaRS structures show a conserved interaction pattern of two arginine residues at renumbered positions 698 and 1786, which were identified in 482 of 524 (92%) structures. The two arginine residues grasp the adenosine phosphate part of the ligand (Fig 4D) with their side chains, resembling a pair of tweezers (Fig 4E), and were thus named Arginine Tweezers. These two arginines are invariant in sequence (Fig 4F). Examples for the Arginine Tweezers motif are residues 217 (corresponding to renumbered residue 698) and 537 (corresponding to renumbered residue 1786) in Class II AspRS structure PDB:1c0a chain A. Additionally, a highly conserved glutamic acid is the most prevalent at renumbered position 700. This residue establishes hydrogen bonds to the adenine group of the ligand in SerRS, HisRS, ThrRS, LysRS, ProRS, and AspRS.

The Backbone Brackets and their counterpart, the Arginine Tweezers, are both responsible for the interaction with the adenosine phosphate part of the ligand (all ligand interactions are shown by example in S2 Fig). Mappings of the motif residues to original sequence numbers can be found in S7 File and S9 File. For some structures it was not possible to pinpoint the conserved motifs after unifying sequence positions (listed in S8 File and S10 File).

Further analysis of secondary structure elements for both motifs shows that residues of the Backbone Brackets are predominantly tied to unordered secondary structure elements (S3 Fig). However, the positions 275, 276, 277, 1359, and 1360 feature a consistently unordered secondary structure. A predominantly unordered state can also be observed for the N-terminal Arginine Tweezers residue 698, while the following three positions almost exclusively occur in strand regions (S4 Fig). Residue 1786 is always observed in *α*-helical regions, mostly at the third position of the *α*-helix element.

The high conservation of backbone or side chain geometry of these motifs suggests that their residues are indispensable for enzyme functionality. To substantiate this assumption, Backbone Brackets and Arginine Tweezers were characterized in greater detail and analyzed regarding their ligand interactions and geometric properties.

### Interaction patterns

Contacts between ligands and proteins are established via a variety of noncovalent interaction types such as hydrogen bonds, *π*-stacking, or salt bridges. These interaction types were annotated using the Protein-Ligand Interaction Profiler (PLIP) [76] to investigate whether evolution adapted entirely different strategies or if some characteristics are shared between both aaRS classes.

Two sets of 29 and 40 representative complexes for Class I and Class II were composed to analyze adenosine phosphate-binding. For the comparison of commonly interacting residues between different aaRS Types, a matrix visualization was designed (Fig 5). This allows for the assessment of interaction preferences at residue level. Data for frequent interactions was available for 12 residues and 10 different aaRS Types for Class I as well as 13 residues and 11 aaRS Types for Class II. All sequence numbers shown in Fig 5 originate from the MSA renumbering and corresponding sequence numbers of all structures in the dataset can be derived from the tables provided in the S13 File and the S14 File.

**Fig 5.**
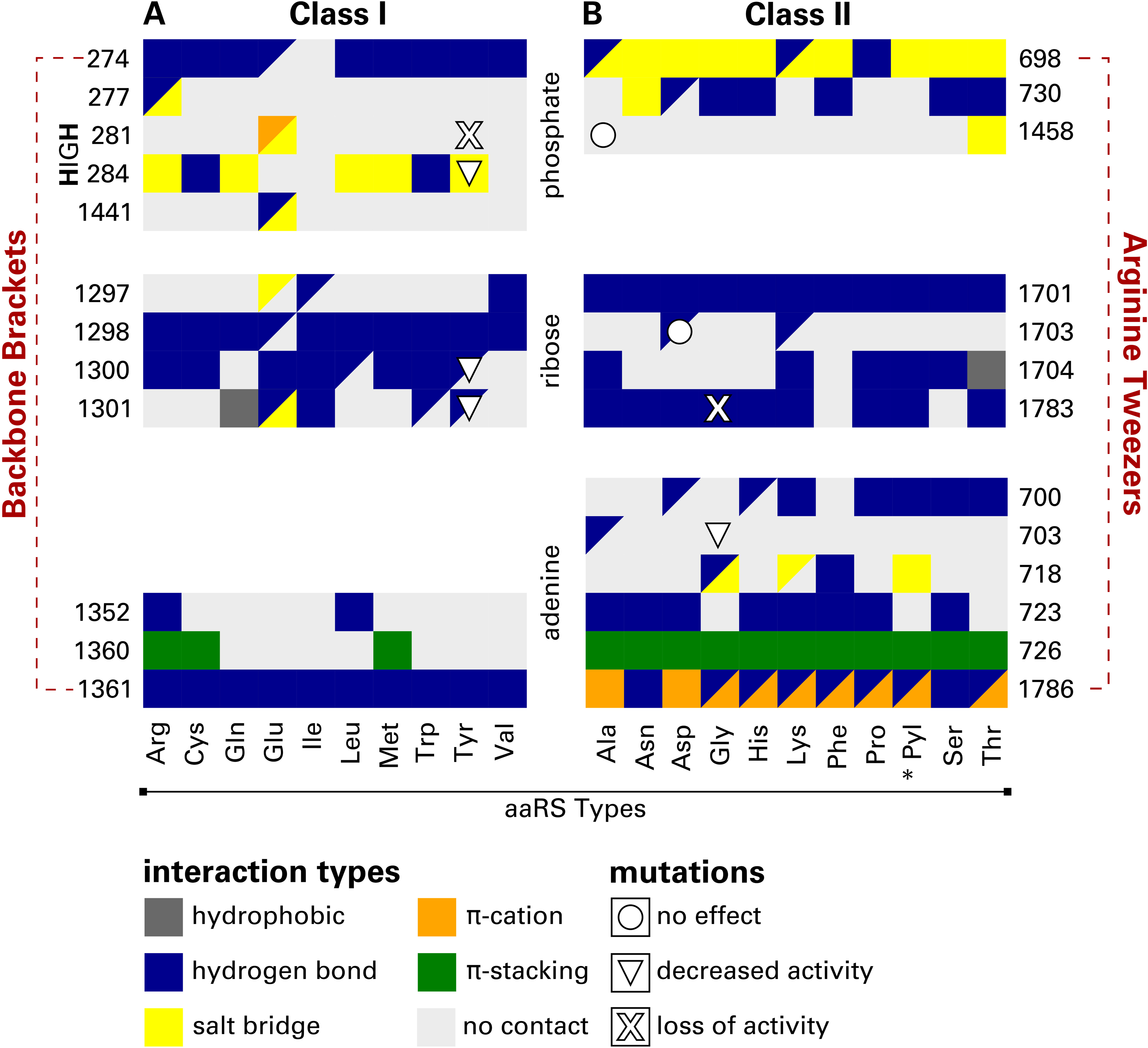
Protein-ligand contacts in representative adenosine phosphate-binding complexes for aaRS Class I and Class II. Residues are grouped according to the non-amino acid ligand fragment (phosphate, ribose, or adenine) that they are interacting with. Preferred interaction types for each aaRS Type and binding site residue are color-coded. Fields split into two triangles indicate two equally preferred interactions. The asterisk (*) indicates aaRS Types incorporating noncanonical amino acids. Automatically retrieved [77, 78] mutation effects [79–85] are shown as centered shapes. In essence, Class I interactions are mainly hydrogen bonds, while Class II adenosine phosphate-binding is realized by an array of different interaction types. All sequence numbers are given according to the MSA.

While six different interaction types are used to bind the adenosine phosphate ligand, hydrogen bonds are the prevalent type of contact, especially for the recognition of the ribose moiety (see Fig 5). The aromatic ring system of adenine is recognized via hydrogen bonds and *π*-stacking interactions in both Class I and Class II complexes.

Class II aaRS bind this part of the ligand also forming *π*-cation interactions with the charge provided by one guanidinium group of the Arginine Tweezers (residue 1786).

Residue 698 interacts predominantly with the negatively charged phosphate group of the ligand via salt bridges. This binding pattern is conserved in Class II and handled by the other guanidinium group featured by the Arginine Tweezers. In Class I, hydrogen bonding is essential for the binding of phosphate. Here, residue 274 binds to the phosphate and is part of the Backbone Brackets motif which embraces the phosphate and the aromatic ring at the other end (residue 1361) using backbone hydrogen bonds.

Both motifs share the tendency to form electrostatic interactions with the *α*-phosphate of the ligand. In general, the phosphate group predominantly participates in salt bridges and hydrogen bonds. The ribose moiety is almost exclusively stabilized by hydrogen bonds to its hydroxyl groups.

### Geometric characterization

Backbone Brackets and Arginine Tweezers were analyzed at the geometrical level to further substantiate the profound differences in adenosine phosphate recognition. The side chains of the Backbone Brackets’ residues are expected to exhibit higher degrees of freedom in comparison to the Arginine Tweezers. Furthermore, a significant change in alpha carbon distance of both motif residues indicates a conformational change during ligand binding. The state complexed with adenosine phosphate (M1) and the state in which no adenosine phosphate is bound (M2) were analyzed separately in order to quantify these aspects (see S1 Fig for a visual representation of M1 and M2). Structure alignments of both motifs in respect to their binding modes (provided in S7 Fig) visually support the differences in side chain orientation and variable amino acid composition of the Backbone Brackets.

The angle between side chains of the Backbone Brackets is continuously high: a mean of 144.90 20.93*^◦^* for M1 and 141.40 20.13*^◦^* for M2, respectively. This emphasizes that the side chain orientation is indistinguishable between M1 and M2 as only the backbone participates in ligand binding. The alpha carbon distance is conserved for the majority of the Backbone Brackets observations, with a mean of 17.92 0.86 Å for M1 and 18.41 0.82 Å for M2, respectively. However, some observations (structures PDB:5v0i chain A, PDB:1jzq chain A, PDB:3tzl chain A, PDB:3ts1 chain A) exhibit higher alpha carbon distances of 20.54 Å, 19.74 Å, 19.10 Å, and 18.79 Å, respectively. In contrast, one occurrence of the Backbone Brackets motif in structure PDB:4aq7 chain A has a remarkably low alpha carbon distance of 16.50 Å. Nevertheless, alpha carbon distances between bound and unbound state differ significantly (*p<*0.01, S5 Fig). This indicates the substantial contribution of backbone interactions as well as the conformational change observed during adenosine phosphate-binding.

The side chain variation is marginal for the Arginine Tweezers if an adenosine phosphate ligand is bound. In contrast, the side chain angle of the apo form is highly variable with a mean of 91.82 8.69*^◦^* for M1 and 79.81 21.67*^◦^* for M2, respectively. The side chain angles between the bound and unbound state differ significantly (*p<*0.01, S6 Fig), reinforcing the pivotal role of highly specific side chain interactions during ligand binding. This effect cannot be observed for the alpha carbon distances of the Arginine Tweezers, with a mean of 14.76 0.66 Å for M1 and 14.93 0.79 Å for M2, respectively.

### Relations to known sequence motifs

Fig 7 encompasses structure and sequence motifs as well as the sequence conservation scores of the underlying MSA. Amino acids interacting with the adenosine phosphate of the ligand (ordinate in Fig 5) are annotated.

For Class I sequence motifs [25, 46, 86], the HIGH motif features sequence conservation and is located nine positions downstream of the N-terminal Backbone Brackets residue. The KMSKS motif exhibits no sequence conservation and can be observed downstream of the C-terminal Backbone Brackets residue. The five-residue motif contains the ligand binding site residue 1441 and is distributed within a corridor of around 70 aligned sequence positions.

For the Class II sequence motifs [25, 57, 58, 60, 86], Motif “1” is moderately conserved in sequence. However, it does not interact with the ligand according to our analysis. Motif “2” is conserved around the N-terminal Arginine Tweezers’ residue and contains five additional ligand binding site residues of lower sequence conservation. Motif “3” exhibits high sequence conservation and includes the C-terminal Arginine Tweezers’ residue.

Further ligand binding site residues, which are not part of known sequence motifs, are mostly occurring in the sequence conserved regions which predominantly bind the ribose moiety.

Fig 7C relates the identified Backbone Brackets and Arginine Tweezers to the proposed Protozyme and Urzyme regions of both aaRS classes [8, 9, 11]. One Backbone Bracket residue is present in the Class I Protozyme, located upstream of the HIGH motif. The other Backbone Bracket residue is located close to the KMSKS motif and therefore part of the Urzyme. Regarding Class II, the N-terminal arginine residue is located in Motif “2” and close to the antisense coding position of the C-terminal Backbone Bracket residue in the Protozyme. The C-terminal Arginine Tweezers residue is located in Motif “3”, that is neither part of the Urzyme nor the Protozyme region.

### Urzyme regions and codon assignment

Rodin and Ohno proposed regions that are associated with each other across the Class division of aaRS [8]. The “HIGH-Motif 2” region was mapped to residues with numbers between 255 to 336 in the renumbered structures of Class I and to 648 to 718 in Class II (according the 46-mers generated by Martinez-Rodriguez et al. [9]). Further, the “KMSKS-Motif 1” region was mapped to residue numbers 1352 to 1452 for Class I and 347 to 371 for Class II in the renumbered structures (according to the alignments by Rodin and Ohno [8]).

Original codons have been mapped for key regions and consensus codons were generated for each of the residues of this region (see Table 1 and Table 2). The codons are rather diverse, but for key positions the middle base exhibits conservation. In the “HIGH-Motif 2” region positions 274-698, 281-692, and 284-689 show complementary middle base pairing. For the “KMSKS-Motif 1” region only one conserved complementary middle base pairing is present at position 1414-365.

**Table 1.**
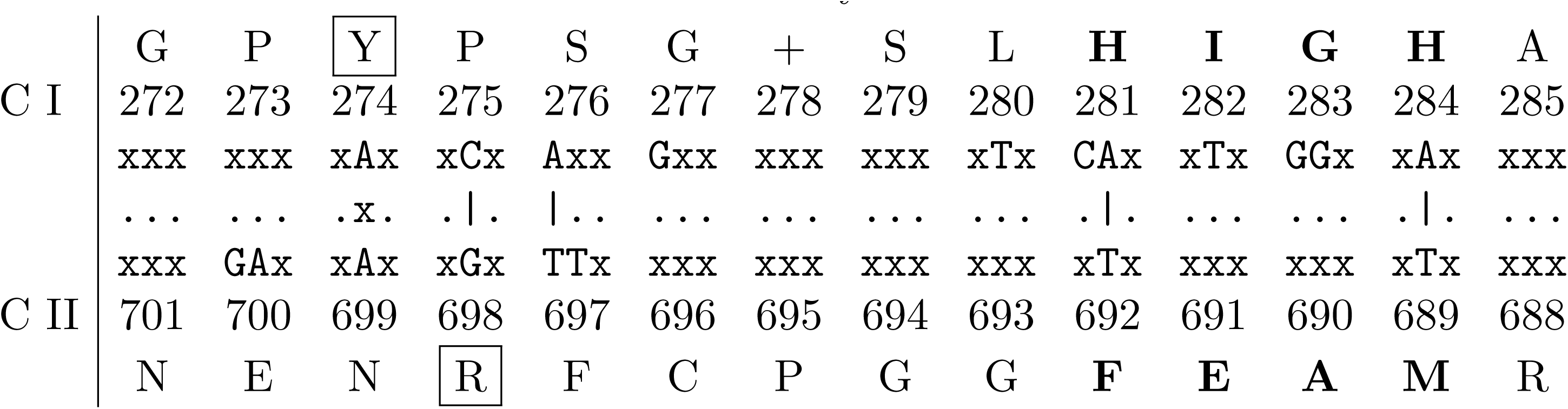
“HIGH-Motif 2” codon assignment and base pairing. First and last row are consensus residues according to the structure-based MSA, “+” indicates gaps. Signature regions according to [8] are emphasized. Sequence numbers are given according to the MSA. Middle rows indicate consensus codons; unassigned positions are indicated by dots, matches by vertical lines, and mismatches by “x”. Arginine Tweezers’ and Backbone Brackets’ residues are framed by boxes.

**Table 2.**
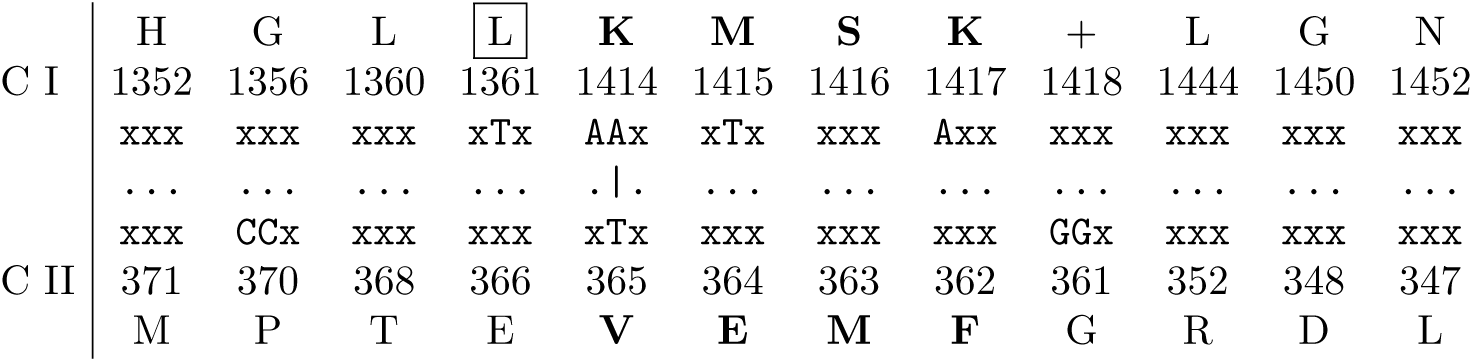
“KMSKS-Motif 1” codon assignment. First and last row are consensus residues according to the structure-based MSA, “+” indicates gaps. Signature regions according to [8] are emphasized. Sequence numbers are given according to the MSA. Middle rows indicate consensus codons; unassigned positions are indicated by dots, matches by vertical lines, and mismatches by “x”. The C-terminal Backbone Brackets’ residue is framed by a box. Sequence positions were omitted if both complementary sequences feature low occupancy, and are therefore not necessarily consecutive.

**Table 3.** Averaged backbone RMSD values of Backbone Brackets and Arginine Tweezers after superimposition.

| motif | binding mode | observations | backbone RMSD [Å] |
| --- | --- | --- | --- |
| Backbone Brackets | M1 | 28 | 0.32 |
| Backbone Brackets | M2 | 59 | 0.34 |
| Arginine Tweezers | M1 | 39 | 0.24 |
| Arginine Tweezers | M2 | 47 | 0.28 |

M1 contains an adenosine phosphate ligand, no adenosine phosphate ligand in M2.

### Effect of mutagenesis experiments and natural variants

To estimate the importance of certain ligand interactions, one can exploit data derived from mutagenesis experiments and natural variants.

Fig 5 shows the effect of nine mutations on the enzymatic activity of aaRS. There is no obvious link between conserved interactions and outcomes of mutations. For example, there are loss of function mutations occurring in regions with observed interactions and equally many cases where no interactions were observed while the mutation still has a negative effect. All sequence positions are given according to the MSA.

For Class I TyrRS, mutations of any histidine of the HIGH motif [46] lead to a decrease in activity, since both residues contribute to the stabilization of the transition state of the reaction [79, 80]. The same holds true for Asp-1300 and Gln-1301 which interact with the ribose part of the ligand [83, 85].

Cys-1458 in Class II AlaRS is part of a four residue zinc-binding motif [88] and an exchange with serine results in no effect whatsoever. It is assumed that the other three amino acids can compensate the mutation [82]. The single-nucleotide polymorphism (SNP) with no known effect is associated to position 1703 in AspRS (rs1803165 in dbSNP [89]).

Ile-703 in Class II GlyRS does not directly interact with the ligand – mutations, however, result in a negative effect and are most prominently linked to Charcot-Marie-Tooth disease as the amino acid is crucial for tRNA ligation [81]. Another SNP occurs at Gly-1783; the exchange with arginine prohibits ligand binding and was tied to a loss of activity as well as distal hereditary motor neuropathy type VA [84].

## Discussion

The reflexive system of building blocks and building machinery implemented in aaRS is an intriguing aspect of the early development of living systems. There is evidence that proteins arose from an ancient set of peptides [90] and that these peptides were co-factors of the early genetic information processing by RNA.

Sequence-based analyses were among the first tools to investigate the transfer of genetic information. DNA and protein sequences comprise the developmental history of organisms, their specialization, and diversification [45]. However, following the “functionalist” principle in biology, sequence is less conserved than structure, which is, in turn, less conserved than function [91]. Therefore, structural features and molecular contacts have been recognized as key aspects in grasping protein function [92, 93] and evolution. Only if the necessary function can be maintained by compatible interaction architectures, the global role of the protein in the complex cellular system is ensured [94]. This is also eminent in aaRS precursor structures that were described to be molten globules but as long as the function of the protein is ensured, it is able to survive during evolution [9]. If evolution tries to conserve structure over function, the evolutionary progress might have been considerably slower and thresholds for the development of new functions would have been higher [91].

Each amino acid of a protein fulfills a certain role and can often be replaced by amino acids with compatible attributes [92]. By considering each amino acid in the context of its sequence, its structural surroundings, and finally its biological function, one can determine possible exchanges and the evolutionary pressure driving these changes [91, 95]. Up to this point, pure sequence or structure analysis methods – ignoring ligand interaction data – missed the functional relevance of the Backbone Brackets entirely.

### Backbone Brackets and Arginine Tweezers

The analysis of Backbone Brackets geometry showed a high variance of side chain angles for both binding modes. The distinction between these modes is significantly manifested in a change of the alpha carbon distance, which supports that the conformational change during ligand binding previously observed in ArgRS [96], TyrRS [47–50, 97], and TrpRS [51, 53–55] is a general mechanism in Class I aaRS. Furthermore, the C-terminal residue of the Backbone Brackets is located close to the KMSKS motif (Table 2). Thus, the structural rearrangement in the KMSKS motif upon ATP binding might indirectly affect the geometric orientation of the C-terminal residue of the Backbone Brackets – especially regarding the position of its alpha carbon.

In contrast to the Backbone Brackets, the Arginine Tweezers are highly restrained in side chain orientation if a ligand is bound, which shows that this orientation is key to adenosine phosphate recognition. If no ligand is bound, the Arginine Tweezers’ geometry is less limited, which is reflected in a higher variability of side chain orientations. Conclusively, the distinction between the two binding modes can be made by taking the geometry of the motifs into account: alpha carbon distances for Backbone Brackets and side chain angles for Arginine Tweezers.

The conserved Arginine Tweezers motif resembles a common interaction pattern for phosphate recognition [92], which usually features positively charged amino acids [98]. However, the conformational space of ATP ligands was shown to be large throughout diverse superfamilies [99] and hence the geometry of binding sites involved in ATP recognition is manifold. The uniqueness of aaRS compared to other ATP-binding proteins was shown in AspRS, where the ligand binds in a compact form with a bent phosphate tail instead of the usually found extended form [99]. This conformation of ATP is energetically unfavorable but allows easy access of the *α*-phosphate for tRNA binding [100]. In general, the nucleophilic attack to the *α*-phosphate of ATP is oppositely directed in Class I and Class II aaRS which possibly evolved at prebiotic time [101]. Quantum mechanical calculations have shown that a lesser propensity for the nucleophilic attack of Class II amino acids is compensated by the bent state of ATP, related binding site residues, and magnesium ions [101]. This specialized mechanism in Class II aaRS suggests that the Arginine Tweezers motif possesses a unique geometry and is not a generalizable pattern for ATP binding, such as the frequently occurring P-loop domain [98].

As the function of fixing the location of the adenosine phosphate part is crucial in aaRS enzymes, mutations of the Arginine Tweezers residues result in loss of function [102, 103]. However, to our knowledge, the Backbone Brackets motif was not identified in earlier literature and is herein described for the first time. The stunning balance of evolutionary diversification [104] and equality in function is underlined by profoundly different implementation of ligand recognition in terms of adjacent sequence (Fig 4C and 4F), embedding secondary structure elements (S3 Fig and S4 Fig), geometrical properties (Fig 6), and interaction characteristics (Fig 5).

**Fig 6.**
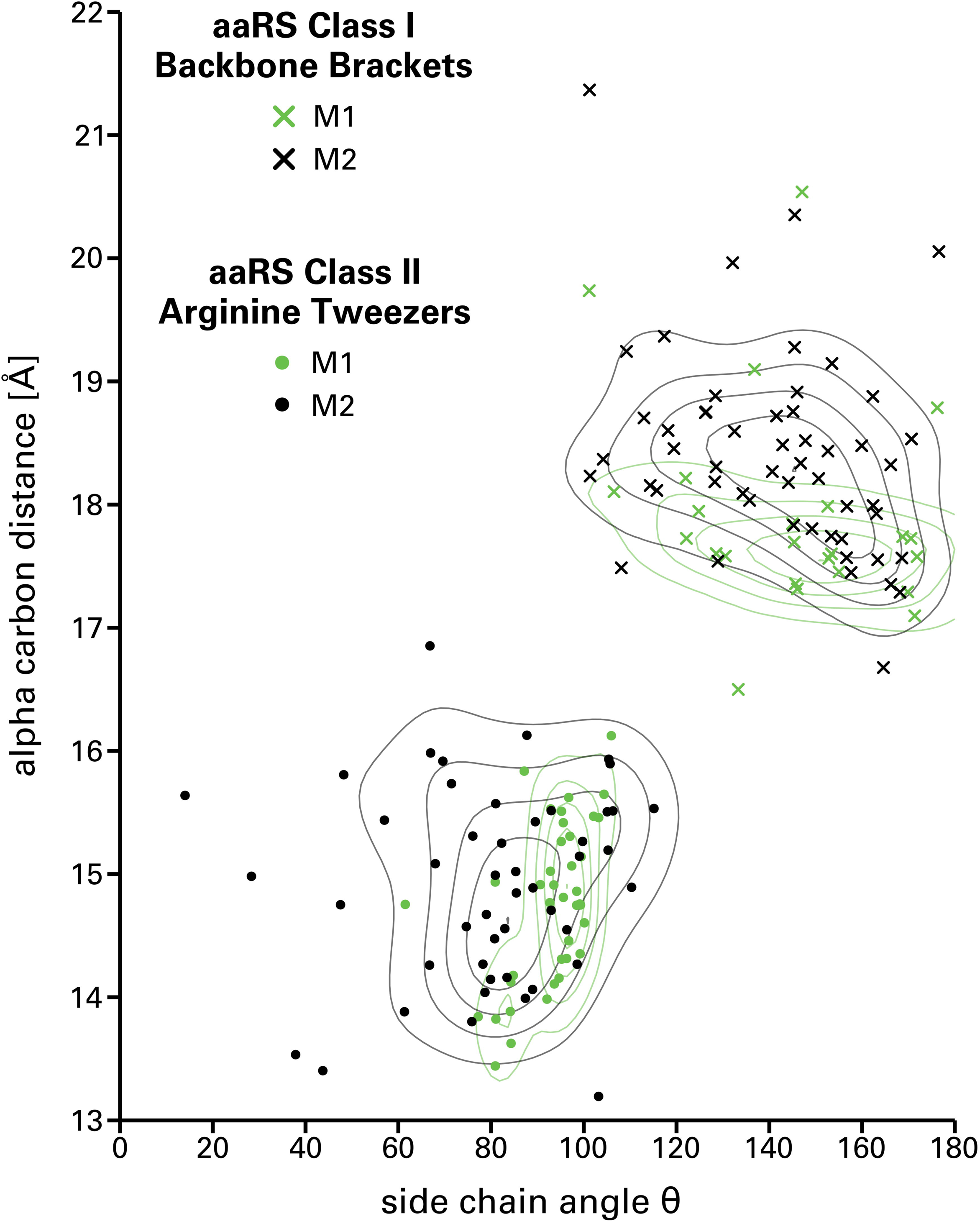
Geometric analysis of the ligand recognition motifs responsible for the adenosine phosphate interaction for aaRS Class I and Class II representative and nonredundant structures. The alpha carbon distance is plotted against the side chain angle θ. Binding modes refer to states containing an adenosine phosphate ligand (M1) or not (M2). Backbone Brackets in M1 allow for minor variance with respect to their alpha carbon distance, constrained by the position of the bound ligand. In contrast, Arginine Tweezers in M1 adapt an orthogonal orientation in order to fixate the ligand.

The catalytic core of both aaRS classes is also hypothesized to consist of amino acids handled by the complementary aaRS Class [11, 105, 106]. The conserved residues of the Arginine Tweezers in Class II support that statement because ArgRS is a Class I aaRS. The contemporary implementations of the Backbone Brackets, however, are dominantly realized by amino acids handled by Class I. Further studies are necessary to test this hypothesis by a detailed investigation of the identified binding site residues for all proteins of the dataset.

### Backbone Brackets are not conserved in sequence

The Backbone Brackets are remarkable, since backbone interactions are often neglected in structural studies. Nevertheless, backbone hydrogen bonds make up at least one-quarter of overall ligand hydrogen bonding [107]. In these cases, side chain properties may only play a minor role, e.g. for steric effects, and allow for larger flexibility in implementation of a binding pattern as long as the correct backbone orientation is ensured. There are examples of protein-ligand complexes where backbone hydrogen bonds are a major part of the binding mechanism, e.g. in binding of the cofactor NAD to a CysG protein from *Salmonella enterica* (PDB:1pjs) as determined with PLIP [76]. In conclusion, the Backbone Brackets exhibit conservation on functional level rather than on sequence level, which renders sequence-based motif analysis infeasible. This motif is a prime example for conservation of function over structure or sequence [91]. When ligands can still be bound specifically by backbone interactions, these binding sites become significantly more resilient to mutations. The complementary codon pairing of both classes’ Protozymes might not only have shaped the genetic code [106], but also required some positions in the Class I Protozyme to be highly variable to compensate changes in the complementary strand. Any amino acid can furnish the observed backbone hydrogen bonds to the ATP ligand, thus drastically increasing the evolvability of both Protozymes.

### Complementary coding of Backbone Brackets and Arginine Tweezers

The isolated “HIGH-Motif 2” region has been shown to be catalytically active [9]. Interestingly, the Arginine Tweezer and the Backbone Bracket appear in very close proximity to each other, when considering the complementary coding according to the Rodin-Ohno hypothesis (see Table 1). This N-terminal Arginine Tweezers residue is oppositely arranged to a conserved proline residue in Class I at position 275. The mapped codons show a matching middle base pair at this position, which is conserved across all kingdoms of life. This further strengthens the evidence for the evolutionary constraints of these residues. Both amino acids fulfill a very important role for the function of aaRS in general. The role of the arginine is well established, it binds the *γ*-phosphate of the ATP molecule and enforces the crucial bent conformation of the phosphate tail [58]. The conserved proline acts as a wedge to open the amino acid binding site to provide access between adjacent strands of a *β*-sheet [32]. The proline residue does not interact directly with the ligand but is still conserved in the binding site, which is why a proposed structural role seems reasonable.

The region reconstructed by [9] is also considered to be the so called Protozyme – the minimal functional aaRS unit required in ancient protein biosynthesis. This region contains the N-terminal residue of both structural motifs identified in this study. This suggests that both N-terminal residues can fulfill their functional role in isolation, but with reduced efficiency. During evolution, the aminoacylation reaction was further improved by adding their other functionally equivalent counterpart.

This is substantiated by the occurrence of the second Backbone Brackets residue at position 1361 very close to the KMSKS mobile loop (residues 1414 to 1417). This C-terminal Backbone Brackets’ residue is part of the region identified as Urzyme, which evolved from the Protozyme, and is more efficient in catalyzing the aminoacylation reaction. Despite the low conservation on sequence, both Backbone Brackets residues have conserved central codon base pairs. This is also the case for other residues that are highly conserved on amino acid sequence, such as the histidine residues in the HIGH motif. This underlines the functionalist principle that has recently been addressed in the context of the evolution of binding sites [91]. The attempt to find conservation on sequence or even structural level is in this case futile, since the interaction is mediated by backbone atoms and in principle this interaction can be realized by any amino acid. Yet, the middle base of the codon for both Backbone Brackets’ residues is conserved. The C-terminal Backbone Brackets’ residue shows a tendency for hydrophobic amino acids (see Fig 4C). This is reflected by the conserved thymine middle base that usually codes for hydrophobic amino acids such as leucine, isoleucine, and valine. In contrast, the conserved adenine middle base of the N-terminal Backbone Brackets’ residue codon encodes for many diverse amino acids, such as glutamic acid, lysine, or glutamine. This coincides with the low sequence conservation observed at this position.

The second Arginine Tweezers residue is situated in the Motif “3” region that has been described before as being important in the aminoacylation reaction [62]. Even though this region is not considered part of either the Ur- or Protozyme, it is present in most of the Class II structures. A comparison of the catalytic rate enhancement, relative to the uncatalyzed second-order rate for the Urzyme with added Motif “3”, but without the preceding insertion domain (similar comparisons have been made previously [45, 108] and are concluded in [9, 11]) is reasonable. It seems that Class II compensated the lack of the second binding element of the ATP part by focusing on the dimerization associated to most of Class II synthetases [45, 109]. In contrast, Class I evolved the C-terminal Backbone Brackets’ residue and did not develop mechanisms such as dimerization to match the reaction speed of Class II. During the course of evolution the ATP binding by two entities proved efficient and was adapted by Class II synthetases as well.

According to the Rodin-Ohno hypothesis [8], one can conclude the following chronological appearance of the Backbone Brackets and Arginine Tweezers motif. The N-terminal residues of both motifs seem to be the most ancient parts, both located in the Protozyme region. Over a prolonged period of time the C-terminal Backbone Brackets residue, which is located close to the KMSKS motif and hence part of the Urzyme, was introduced. The most recent residue seems to be the C-terminal Arginine Tweezers’ residue, located in Motif “3”, which is neither part of the Protozyme nor the Urzyme.

### Disease implications

Due to the fundamental role of aaRS for protein biosynthesis, a systematic assessment of mutation effects in yeast was conducted by Cavarelli and coworkers [102]. Mutations of aaRS-coding genes can be drastic and may result in a variety of human diseases, even if the structural effect is unknown [110, 111].

Structural analysis of a GlyRS mutant (G526R) showed that the Charcot-Marie-Tooth disease may be caused by blockage of the ATP binding site. Furthermore, this mutation induces a larger contact area in the homo-dimer interface, which stems partially from the anticodon binding domain [84]. Other mutations result in a wider range of diseases and symptoms such as hearing loss, ovarian failure, or cardiomyopathy [112, 113]. Even for cellular processes unrelated to translation, aaRS play a pivotal role, e.g. for angiogenesis [114]. Due to the highly individual characteristics of aaRS enzymes between organisms, it is possible to create precisely targeted antibiotics with minimal side effects [115–117].

Unfortunately, automatically mapped mutational data does not cover the Backbone Brackets or Arginine Tweezers motif. It is expected that mutations of the Arginine Tweezers will cause a strong decrease in enzyme activity as shown in [102]. In contrast, the Backbone Brackets are expected to be more resilient to mutational events. However, bridging the gap between mutational studies and key interaction patterns will require further analysis beyond this study and needs to be substantiated by *in vitro* experiments. The provided high-quality aaRS dataset can serve as the basis for such work.

## Limitations

The method used to unify residue numbering in all structures relies on the quality of the used MSA as well as the quality of local structure regions. Hence, the Backbone Brackets and Arginine Tweezers were not successfully mapped for all structures of the dataset. On the one hand, some binding site regions were not experimentally determined (e.g. PDB:3hri) or the mapping of the motif residues failed (e.g. PDB:4yrc) due to ambivalent regions in the MSA. On the other hand, some aaRS may have evolved different strategies to bind the ligand, even for the same aaRS type [118].

However, the conserved ligand interactions were related to known sequence motifs (Fig 7). The sequentially high variance of the KMSKS motif was described before [46] and explains why the MSA algorithm distributes this motif over 70 positions. Another explanation is the differing conformation between the two binding modes [47–49, 51–55] which leads to a scattered structure-based sequence alignment in the KMSKS region [74]. The interacting residues 1352, 1360, and 1361 of Class I are located upstream of the KMSKS motif. In case of Class I, the AIDQ motif in TrpRS is known [110], yet no consensus for all aaRS Types was established. Class II sequence motifs exhibit high degeneracy and can hardly be identified without structural information [104]. Motif “1” is the only sequence motif which is not linked to any relevant ligand interaction site; its primary role lies in the stabilization of Class II dimers [57].

**Fig 7.**
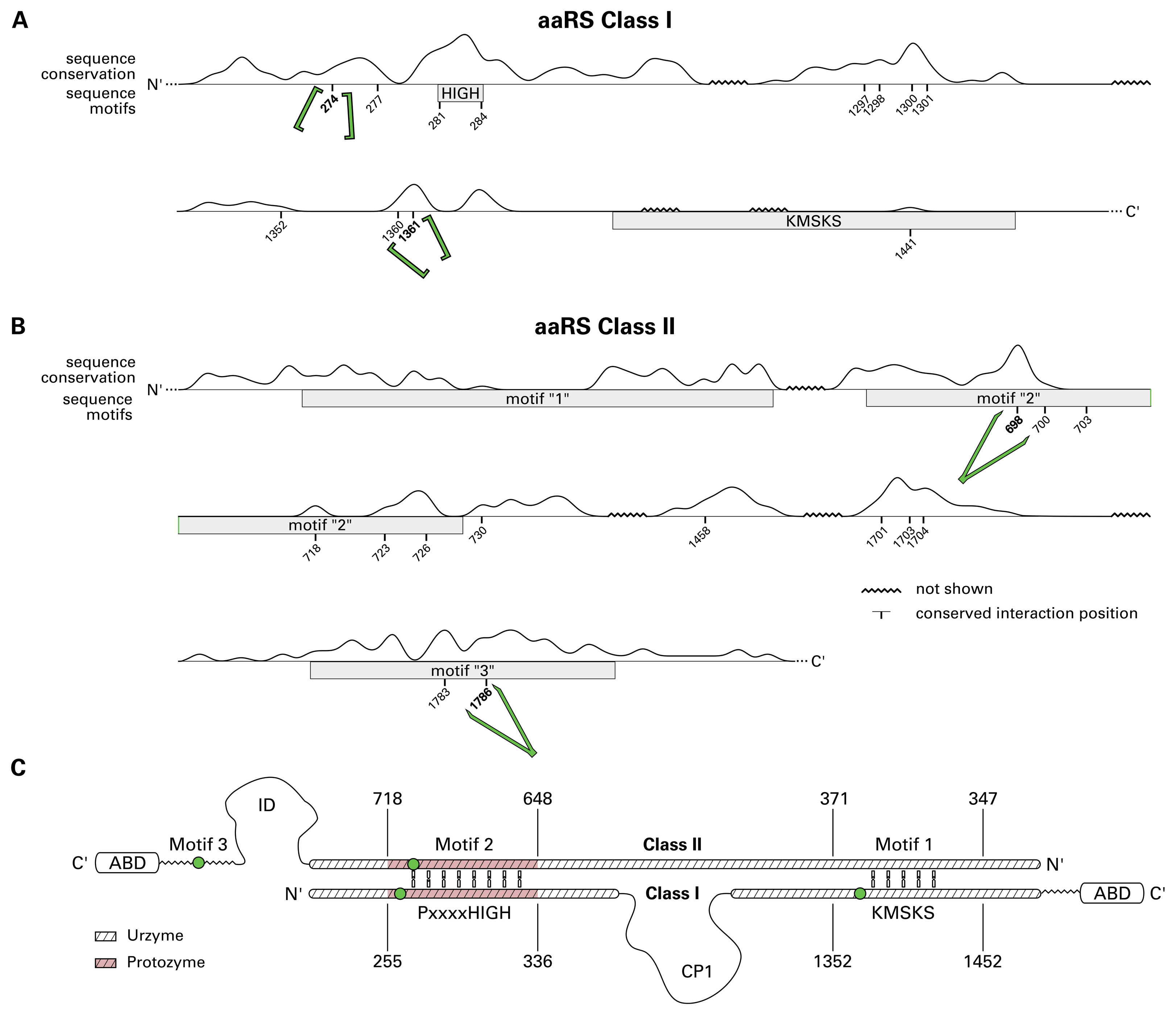
Integrative sequence view for aaRS Class I (A) and Class II (B). Boxes delineate sequence motifs previously described in literature [46, 57, 58]. The trace depicts the sequence conservation score of each position in the MSA (S5 File and S6 File). These scores were computed with Jalview [40, 87], positions composed of sets of amino acids with similar characteristics result in high values. Furthermore, all positions relevant for ligand binding (Fig 5) are depicted. Backbone Brackets and Arginine Tweezers have been emphasized by their respective pictograms. Positions of low conservation or those not encompassed by sequence motifs were intangible to studies primarily based on sequence data. Especially backbone interactions might be conserved independently from sequence. (C) Sequence representation of the Rodin-Ohno hypothesis [8, 9, 11] with equivalents of the Backbone Brackets or Arginine Tweezers residues shown as green dots. The N-terminal residue of each, the Backbone Brackets and the Arginine Tweezers motif, is present in the Protozyme region (shaded red). Additionally, the C-terminal Backbone Brackets’ residue is located in the Urzyme region.

The geometric characterization of the two ligand recognition motifs (see Fig 6) highlighted some observations of the Backbone Brackets, which exhibit a substantial increase or decrease of the residue alpha carbon distance. For instance, chain A of an LeuRS of *Escherichia coli* (PDB:4aq7) is complexed with tRNA and the Backbone Brackets’ alpha carbon distance is about 1 Å below the average. Manual investigation of this structure showed that there is no obvious conformational difference to other structures. Likewise, the annotated interactions were checked for consistency using PLIP and showed usual interactions with the adenine and the sulfamate group (the phosphate analogue) of the ligand. For the Backbone Brackets with higher alpha carbon extent (structures of IleRS, TrpRS, and TyrRS), interaction analysis revealed that residue 274 interacts with the amino acid side chain, as all of these structures contain a single aminoacyl ligand (PDB:3tzl chain A, PDB:3ts1 chain A, PDB:1jzq chain A) or two separate ligands (amino acid and AMP, PDB:5v0i chain A). This suggests that the structures resemble a partially changed conformation prior to tRNA ligation and a possible role of the Backbone Brackets motif in amino acid recognition. Likewise, these effects can arise from low quality electron density maps in the structure regions of interest. However, these hypotheses have to be addressed and validated in future work.

Interestingly, our analysis did not reveal a high count or conservation of interactions established with the well-known HIGH motif in Class I. Despite irregularly occurring salt bridges, hydrogen bonds, and one *π*-cation interaction in GluRS (see Fig 5A), no interactions were observed. This especially holds true for the first histidine residue of the HIGH motif, which only interacts with the ligand in GluRS. However, it was shown that the HIGH motif is mainly relevant for binding in the pre-acylation transition state of the reaction [46], i.e. HIGH interacts with the phosphate of ATP. This explains the irregular observations of interactions which are established only if an ATP ligand is present (e.g. GluRS PDB:1j09 chain A residue 15).

### Prospects

Adaptations of the presented workflow to other protein families of interest might allow to study binding mechanisms in a new level of detail and by using publicly available data alone. Even if the geometric characterization is dependent on the quality of local structure regions, the comparison of alpha carbon distances and side chain angles is a simple yet valuable tool to separate different binding states. Geometrical properties can reveal the importance of conserved side chain orientations, the degree of freedom in unbound state, or shifts in backbone arrangement. However, choosing these two properties to compare residue binding motifs depends on the specific use case.

For the analysis of aaRS structures, the geometric characterization of the two conserved core interaction patterns was shown to be sufficiently sensitive to suggest the structural rearrangement of Class I aaRS to be a general mechanism. Hence, if structural motifs conserved in a larger number of protein structures are known, geometric analysis can reveal insights into global structural effects that occur during ligand binding without requiring any additional information.

In a similar way, the obtained interaction data proved as a valuable resource to understand fundamental aspects of aaRS ligand recognition. Despite the fact that interactions can not be determined for apo structures and do not take into consideration the dynamic nature of enzyme reactions, both, structure and interaction data conflates several aspects of evolution and proved to outperform pure sequence-based methods. Regarding the Rodin-Ohno hypothesis, structural investigation of the proposed Protozymes [8, 9, 11] and their ligand binding properties can further substantiate the importance of the Backbone Brackets and Arginine Tweezers as the primordial ATP binding site.

The designed approach was used to analyze aaRS from the different viewpoints: sequence backed by structure information, ligand interactions, and geometric characterization of essential ligand binding patterns. Additionally, this study provides the largest manually curated dataset of aaRS structures including ligand information available to date. This can serve as foundation for further research on the essential mechanisms controlling the molecular information machinery, e.g. investigate the effect and disease implications of mutations on crucial binding site residues. Further phylogenetic analyses can be conducted, based on the identified structural motifs. The sequence of aaRS proteins was shown to be highly variable [61] yet Backbone Brackets and Arginine Tweezers constituted a common pattern shared by almost all structures of the corresponding aaRS classes.

Alongside the aaRS-specific results, the workflow is a general tool for identification of significant ligand binding patterns and the geometrical characterization of such. Further studies may adapt the presented methodology to study common mechanisms in highly variable implementations of ligand binding, i.e. for nonribosomal peptide synthetases as another enzyme family that is required to recognize all 20 amino acids [119].

## Materials and methods

### Dataset preparation

Proteins with domains annotated to belong to aaRS families according to Pfam 31.0 [120] were selected (see S1 Appendix for a detailed list of Pfam identifiers) and their structures were retrieved from PDB. Additionally, structures with Enzyme Commission (EC) number 6.1.1.- were considered and included in the initial dataset. Structures with putative aaRS function were excluded.

For each catalytic chain the aaRS Class and Type, resolution, mutational status, the taxonomy identifier of the organism of origin, and its superkingdom were determined. For chains where a ligand was present, these ligands were added to the dataset and it was decided if this ligand is either relevant for amino acid recognition (i.e. contains an amino acid or a close derivate as substructure), for adenosine phosphate-binding (i.e. contains an adenosine phosphate substructure), or for both (e.g. aminoacyl-AMP).

As the presented study focuses on the binding of the adenosine phosphate moiety, two binding modes referred to as M1 and M2 (S1 Fig) were defined. M1 features an adenosine phosphate-containing ligand (e.g. aminoacyl-AMP, ATP), whereas M2 does not contain any ligand that binds to the adenosine phosphate recognition region of the binding pocket (e.g. plain amino acid, empty pocket).

To avoid the use of highly redundant structures for analysis, all structures in the dataset were clustered according to *>*95% sequence identity using Needleman-Wunsch [121] alignments and single-linkage clustering. For each of these clusters, a representative chain (selection scheme listed in S2 Appendix) was determined. The same procedure was used to define representative chains for the adenosine phosphate bound state M1 and no adenosine phosphate bound state M2. The final dataset is provided as formatted table in S1 File and as machine-readable JSON version in S2 File.

### Mapping of binding sites

To allow a unified mapping of aaRS binding sites, an MSA of 81 (75) representative wild type sequences of Class I (Class II) (S3 File and S4 File) aaRS was performed. The alignment was calculated with the T-Coffee expresso pipeline [74], which guides the alignment by structural information. Using the obtained MSA (S5 File and S6 File), residues in all aaRS structures were renumbered with the custom script “MSA PDB Renumber”, available under open-source license (MIT) at github.com/vjhaupt. All renumbered structures are provided in PDB file format (S11 File and S12 File). Only protein residues were renumbered, while chain identifiers and residue numbers of ligands were left unmodified. Lists of structures where the Backbone Brackets or Arginine Tweezers were not mapped successfully are found in S8 File and S10 File.

### Annotation of noncovalent interactions

Annotation of noncovalent interactions between an aaRS protein and its bound ligand(s) was performed with the PLIP [76] command line tool v1.3.3 on all renumbered structures with default settings. The renumbered sequence positions of all residues observed to be in contact with the ligand were extracted. This resulting set of interacting residues was used to determine the position-identical residues from all aaRS structures in the dataset, even if no ligand is bound.

### Generation of interaction matrix

Information on noncovalent protein-ligand interactions from renumbered structure files (see above) was used to prepare separate interaction matrices for aaRS Class I and Class 1. II. First, only representative structures for M1 were selected. Second, only residues which are in contact with the non-amino acid part of the ligand (i.e. adenine, ribose moiety or the phosphate group) were considered. This was validated manually for each residue. Furthermore, residues relevant for only one aaRS Type were discarded. For each considered residue, the absolute frequency of observed ligand interactions was determined with respect to the PLIP interaction types (hydrophobic contacts, hydrogen bonds, salt bridges, *π*-stacking, and *π*-cation interactions). Additionally, the count of residues not interacting with any ligand (“no contact”) was determined. In the interaction matrix (Fig 5), aaRS Types are placed on the abscissa and renumbered residue positions on the ordinate. The preferred interaction type for each residue and ligand species is color-coded. If two interaction types occurred with the same frequency, a dual coloring was used. Residues were grouped in the figure according to the ligand fragment they are mainly forming interactions with.

### Annotation of mutagenesis sites and natural variants

For each chain, a mapping to UniProt [77] was performed using the SIFTS project [78]. Where available, mutation and natural variants data was retrieved for all binding site residues from the UniProt [77] database. In total, 32 mutagenesis sites and 8 natural variants were retrieved.

### Analysis of core-interaction patterns

All motif occurrences in M1 and M2 representative chains were aligned in respect to their backbone atoms (S7 Fig) using the Fit3D algorithm [122]. Additionally, the alpha carbon distances and the angle between side chains were determined. The side chain angle *θ* between two residues was calculated by abstracting each side chain as a vector between alpha carbon and the most distant carbon side chain atom. If *θ* = 0*^◦^* or *θ* = 180*^◦^* the side chains are oriented in a parallel way. Side chain angles were not calculated if one or both residues of the Backbone Brackets motif were glycine.

Furthermore, the sequential neighbors of the core-interaction patterns have been visualized with WebLogo graphics [75], regarding their sequence and secondary structure elements. Secondary structure elements were assigned according to the rule set of DSSP [123].

### Codon assignment

The sequence regions proposed by Rodin and Ohno [8] were chosen as candidates for the codon assignment; the tangible positions are listed in Table 1 and Table 2. Cluster representative structures where chosen for the following analysis. In order to assign the original coding nucleotide sequence to each of the structures, the sequences of the structures were retrieved from the UniProt database [77] using the SIFTS project [78] to map PDB structures to UniProt entries. Afterwards, the corresponding codons were assigned to each amino acid by extracting them from the annotated coding sequences deposited in the European Nucleotide Archive [124]. Consensus codons were generated for each amino acid using WebLogo graphics [75] and choosing the most prominent nucleotide for positions with an entropy higher than one bit.

## Acknowledgments

We thank Peter R. Wills for initially approaching us with the intriguing topic of the origin of genetic coding. Further, we appreciated the meetings and are grateful for guiding us the way through the entire project. Gratitude is owed to Lauren Adelmann, Hanna Siewerts, and Alexander Eisold for proofreading the manuscript. We thank the reviewers who proposed valuable improvements to the manuscript and helped us to strengthen the discussion substantially.

## Supporting information

**S1 Fig. Binding mode definition.** Binding modes M1 and M2 are defined based on the complexed ligand: ligands that bind to the adenosine phosphate moiety (highlighted in red, only in contact when adenosine phosphate is part of the ligand) of the binding site (M1), no ligands or ligands that bind exclusively to the aminoacyl part (green) of the binding site (M2).

**S2 Fig. Core-interaction patterns.** Both aaRS classes contain highly conserved patterns, responsible for proper binding of the adenosine phosphate part of the ligand. Class I aaRS share a highly conserved set of backbone hydrogen interactions with the ligand: the Backbone Brackets. Class II active sites contain a pattern of two arginine residues grasping the adenosine phosphate ligand: the Arginine Tweezers. Interactions were calculated with PLIP [76] and are represented with colored (dashed) lines: hydrogen bonds (solid, blue), *π*-stacking interactions (dashed, green), *π*-cation interactions (dashed, orange), salt bridges (dashed, yellow), metal complexes (dashed, purple), and hydrophobic contacts (dashed grey). (**A**) Class I Backbone Brackets motif and interactions with the ligand Tryptophanyl-5’AMP as observed in TrpRS structure PDB:1r6u chain A. (**B**) Class II Arginine Tweezers motif and interactions with the ligand Lysyl-5’AMP as observed in LysRS structure PDB:1e1t chain A.

**S3 Fig. Secondary structure of Backbone Brackets adjacent residues.** WebLogo [75] representation of secondary structure elements around the Backbone Brackets residues (274 and 1361) annotated by DSSP [123]: helices (blues), strands (red), and unordered (black). Unassigned states are represented by the character ’C’. The height of each character corresponds to the relative frequency.

**S4 Fig. Secondary structure of Arginine Tweezers adjacent residues.** WebLogo [75] representation of secondary structure elements around the Arginine Tweezers residues (698 and 1786) annotated by DSSP [123]: helices (blues), strands (red), and unordered (black). Unassigned states are represented by the letter ’C’. The height of each character corresponds to the relative frequency.

**S5 Fig. Distributions of alpha carbon distances for Backbone Brackets and Arginine Tweezers.** Distributions of alpha carbon distances for Class I Backbone Brackets motif and Class II Arginine Tweezers motif in adenosine phosphate bound (M1) and unbound state (M2). The alpha carbon distance of the Backbone Brackets differs significantly between the two states (Mann-Whitney U *p<*0.01).

**S6 Fig. Distributions of side chain angles for Backbone Brackets and Arginine Tweezers.** Distributions of side chain angle *θ* for Class I Backbone Brackets motif and Class II Arginine Tweezers motif in adenosine phosphate bound (M1) and unbound state (M2). The side chain angles of the Arginine Tweezers differs differs significantly between the two states (Mann-Whitney U *p<*0.01).

**S7 Fig. Alignments of Backbone Brackets and Arginine Tweezers.** Structural backbone-only alignments computed with Fit3D [122] of relevant binding site residue motifs derived from M1 and M2 representative structures in respect to their binding modes for aaRS Class I and Class II. (**A**) The Class I Backbone Brackets motif aligned in respect to binding modes. A high side chain variance (gray line representation) is evident. However, backbone orientations are highly conserved to realize consistent hydrogen bond interaction with the adenosine phosphate part of the ligand. (**B**) The Class II Arginine Tweezers motif aligned in respect to binding modes. Low side chain variance can be observed if an adenosine phosphate ligand is bound (M1), whereas the absence of an adenosine phosphate ligand (M2) allows an increased degree of freedom for side chain movement. Averaged backbone and side chain RMSD values after all-vs-all superimposition are shown in S1 Table.

**S8 Fig. Pairwise sequence and structure similarity.** Structure and sequence similarity for pairs of cluster representative chains for aaRS Class I (**A**) and II (**B**). Depicted is the sequence similarity (% identity) after a global Needleman-Wunsch [121] alignment of both structures against the structure similarity determined by TMAlign [73]. For Class I (Class II) 95% of all pairs exhibit *<*33% (29%) sequence identity and *<*0.85 (0.84) TM score. The 95% quantile borders are depicted as red dashed lines.

**S9 Fig. Origin organisms of aaRS Class I and Class II structures in the dataset.** The organisms of origin for aaRS Class I (**A**) and Class II (**B**) structures in the dataset. The inner circles correspond to the superkingdom of the organism. The outer circle depicts the partition into specific species (combining different strains).

Sections representing eukaryotic species are colored in violet, bacteria are colored in green, archaea are colored in orange and vira are colored in gray. Species, that are origin of less than two percent of the structures are condensed to the “other” cluster for each superkingdom. All superkingdoms are represented in both datasets. Class I contains more bacterial structures than Class II, but fewer originating from eukaryotes or archaea. Interestingly, Class I also contains one viral structure. The Class I set contains four mitochondrial structures, whereas Class II contains 15 mitochondrial structures. Despite the diverse origins of the structures the conserved interaction patterns can be observed.

**S1 Table. Backbone and side chain RMSD of Backbone Brackets and Arginine Tweezers after superimposition.** Averaged backbone and side chain RMSD values after all-vs-all superimposition are shown in this table.

**S1 Appendix. Dataset preparation**

All selected protein chains from the PDB carry one of the following protein family annotations, according to Pfam [120]: PF00133, PF00152, PF00579, PF00587, PF00749, PF00750, PF01406, PF01409, PF01411, PF02091, PF02403, PF02912, PF03485, PF09334. Additionally, structures annotated with an EC number indicating tRNA-ligation activity were selected: 6.1.1.1 (TyrRS), 6.1.1.2 (TrpRS), 6.1.1.3 (ThrRS), 6.1.1.4 (LeuRS), 6.1.1.5 (IleRS), 6.1.1.6 (LysRS), 6.1.1.7 (AlaRS), 6.1.1.9 (ValRS), 6.1.1.10 (MetRS), 6.1.1.11 (SerRS), 6.1.1.14 (GlyRS), 6.1.1.15 (ProRS), 6.1.1.16 (CysRS), 6.1.1.17 (GluRS), 6.1.1.18 (GlnRS), 6.1.1.19 (ArgRS), 6.1.1.20 (PheRS), 6.1.1.21 (HisRS), 6.1.1.22 (AsnRS), 6.1.1.23 (AspRS), 6.1.1.26 (PylRS).

For each of the resulting structures, the existence of a catalytic domain was checked manually and only the chains containing a domain with confirmed catalytic activity were retained. If there were ligands present in the structure, these ligands were annotated manually to avoid errors in the assignment of ligands to their catalytic chain. This procedure resulted in a high quality dataset of 972 individual aaRS chains containing a catalytic domain.

**S2 Appendix. Selection of representative entries**

In order to avoid redundancy, representatives were defined for each sequence cluster with *>*95% sequence identity and discriminated between three types: cluster representatives, representatives that contain an adenosine phosphate ligand (M1), and representatives that do not contain an adenosine phosphate ligand (M2).

The selection criteria for these categories were defined as follows:

- cluster representative:

1. protein must be wild type (if wild type exists)
2. best resolution
3. longest sequence coverage
- representatives with an adenosine-relevant ligand

1. chain must contain an adenosine phosphate ligand
2. this ligand must be standard (adenosine phosphate or close derivate)
3. no experimentally validated inhibitor ligand in the binding site
4. protein must be wild type (if wild type exists)
5. best resolution
- representatives without an adenosine phosphate ligand

1. chain must not contain an adenosine phosphate ligand
2. binding site must not contain an inhibitor ligand
3. protein must be wild type (if wild type exists)
4. best resolution

For ties in the selection, structures were sorted naturally ascending according to their PDB identifier and chain identifier and the first was chosen.

**S1 File. Dataset as table.** Summary table of all aaRS protein chains used for the analysis, including PDB identifier, chain identifier, superkingdom, taxonomy identifier, and ligand information (if any).

**S2 File. Dataset as JSON file.** Machine-readable JSON version of the dataset. Additionally enriched with protein sequence, sequence cluster identifier, and representative types for each dataset entry.

**S3 File. Class I sequences in FASTA format.** Protein sequences of Class I aaRS structures used to construct the structure-guided MSA in FASTA format.

**S4 File. Class II sequences in FASTA format.** Protein sequences of Class II aaRS structures used to construct the structure-guided MSA in FASTA format.

**S5 File. Class I multiple sequence alignment.** Structure-guided MSA of Class I sequences in FASTA format.

**S6 File. Class II multiple sequence alignment.** Structure-guided MSA of Class II sequences in FASTA format.

**S7 File. Backbone Brackets residue mapping.** Mapping of the Backbone Brackets Class I motif to sequence positions in origin structures.

**S8 File. Backbone Brackets failed mapping.** List of structures where the mapping of the Backbone Brackets motif was not possible.

**S9 File. Arginine Tweezers residue mapping.** Mapping of the Arginine Tweezers Class II motif to sequence positions in origin structures.

**S10 File. Arginine Tweezers failed mapping.** List of structures where the mapping of the Arginine Tweezers motif was not possible.

**S11 File. Archive containing Class I renumbered structures.** All structures of Class I aaRS with residues renumbered according to the MSA.

**S12 File. Archive containing Class II renumbered structures.** All structures of Class II aaRS with residues renumbered according to the MSA.

**S13 File. Renumbering table for Class I structures.** Formatted table that contains all sequence positions of the Class I MSA and annotations of sequence motifs, Backbone Brackets’ residues, and ligand binding regions (rows). Each renumbered sequence position is related to its original sequence position for every structure in the dataset (columns).

**S14 File. Renumbering table for Class II structures.** Formatted table that contains all sequence positions of the Class II MSA and annotations of sequence motifs, Arginine Tweezers’ residues, and ligand binding regions (rows). Each renumbered sequence position is related to its original sequence position for every structure in the dataset (columns).

