## Supplementary figures and images for "Backbone Brackets and Arginine Tweezers delineate Class I and Class II aminoacyl tRNA synthetases"

### Supplementary Materials

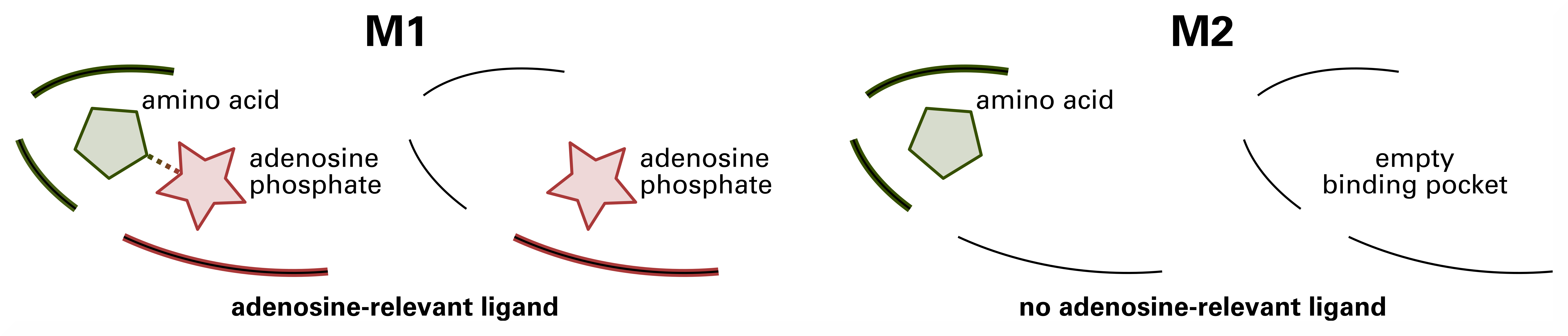

### Supplementary Materials

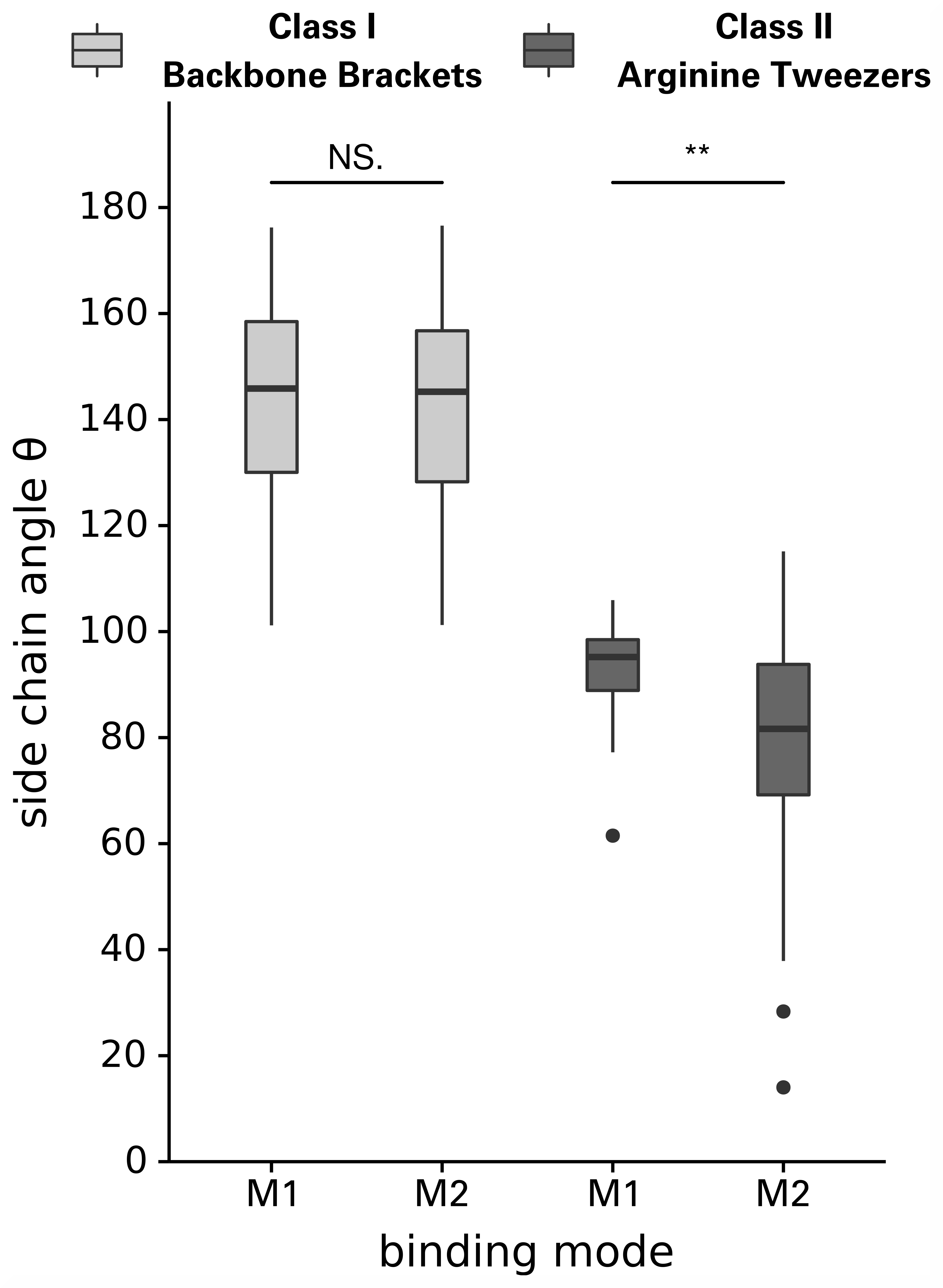

### Supplementary Materials

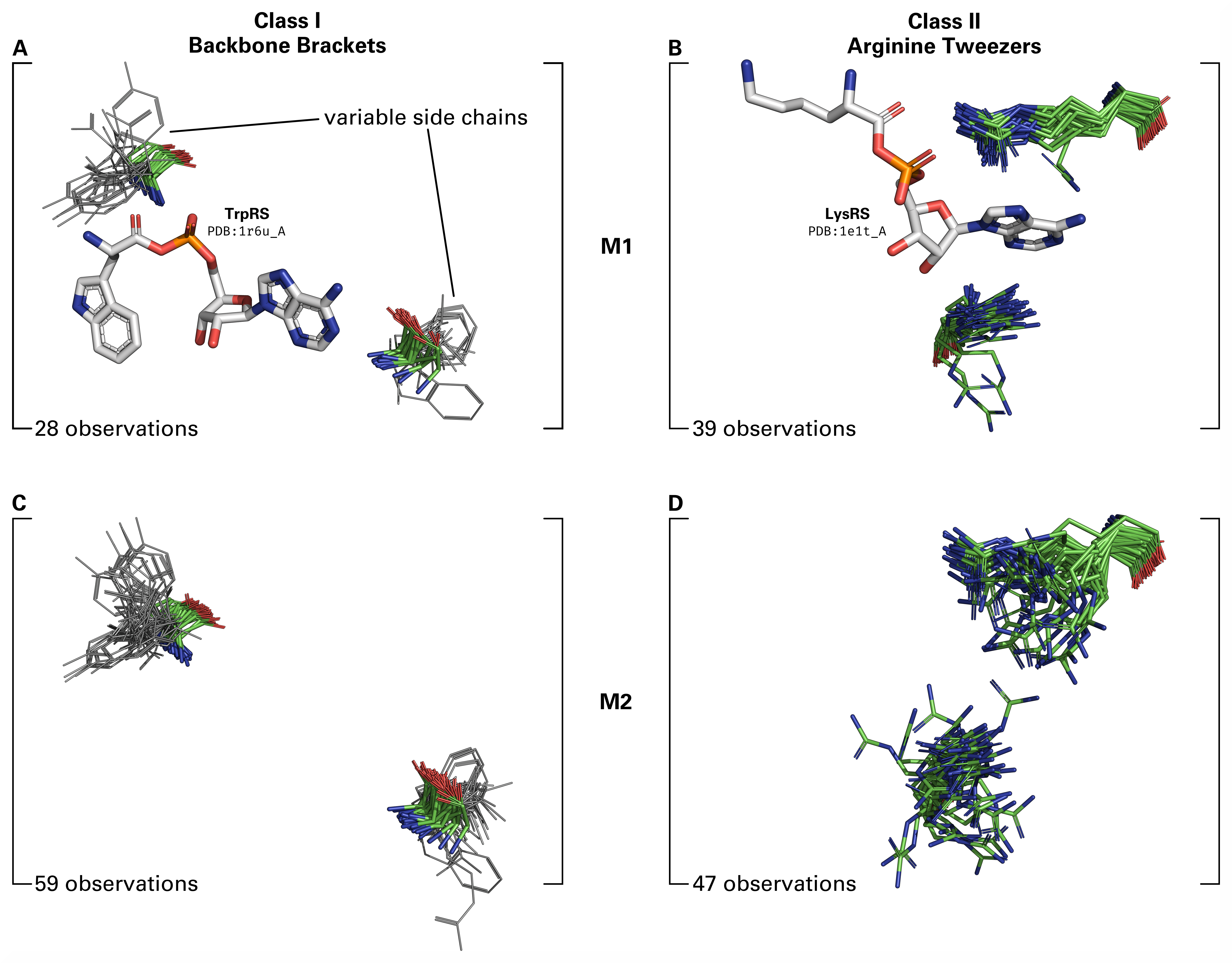

### Supplementary Materials

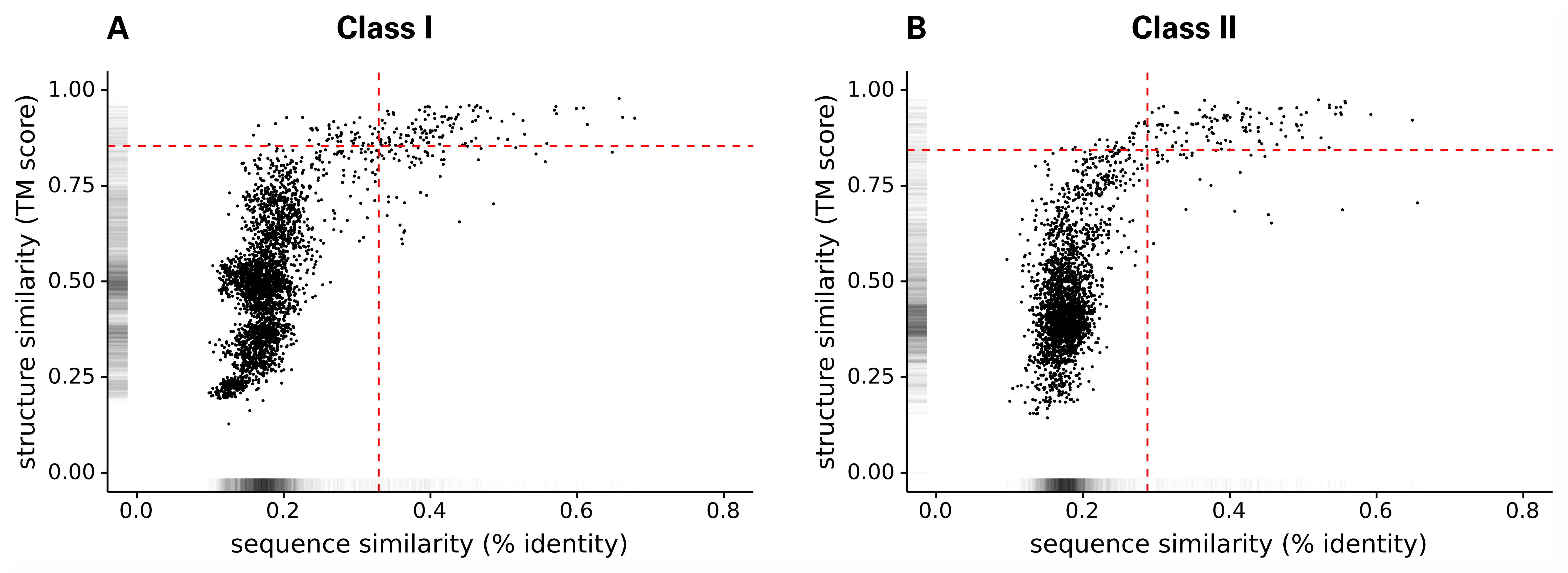

### Supplementary Materials

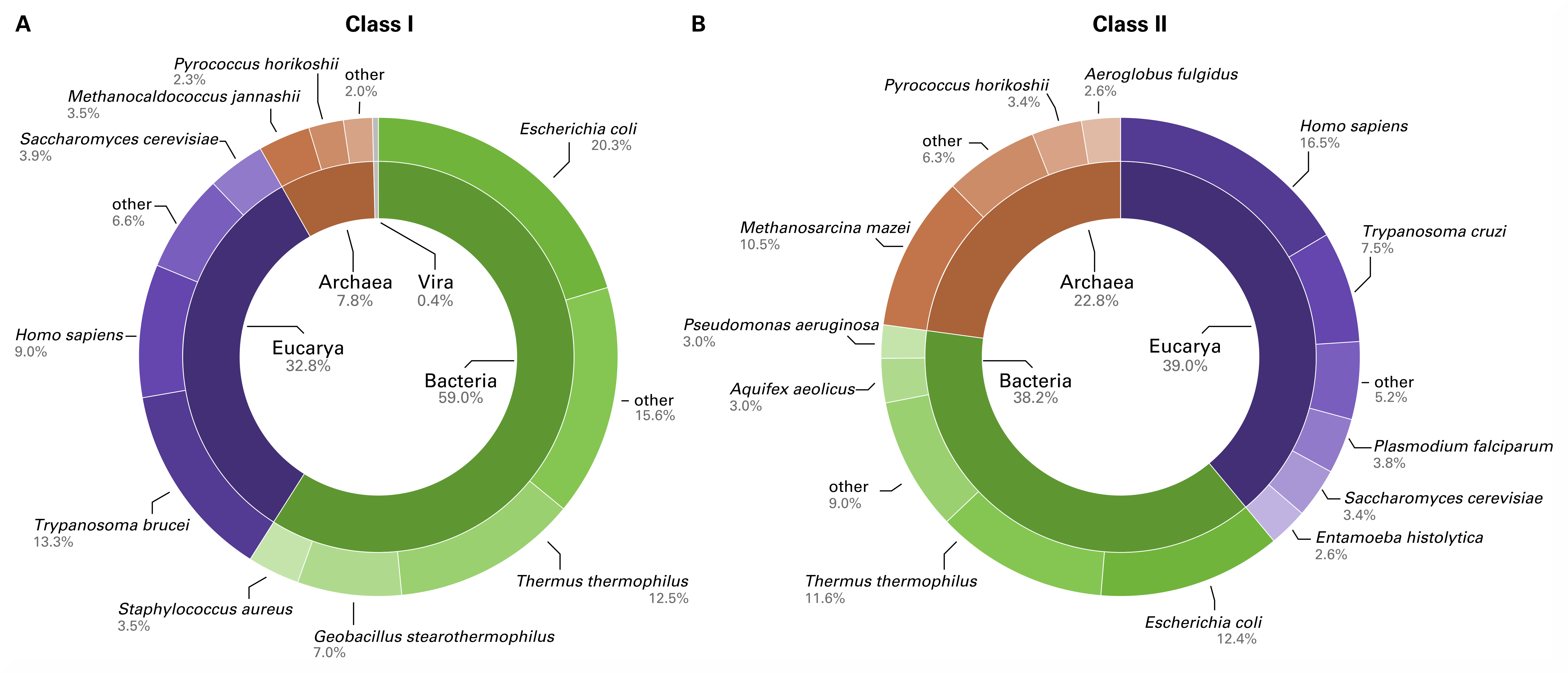

### Supplementary Materials

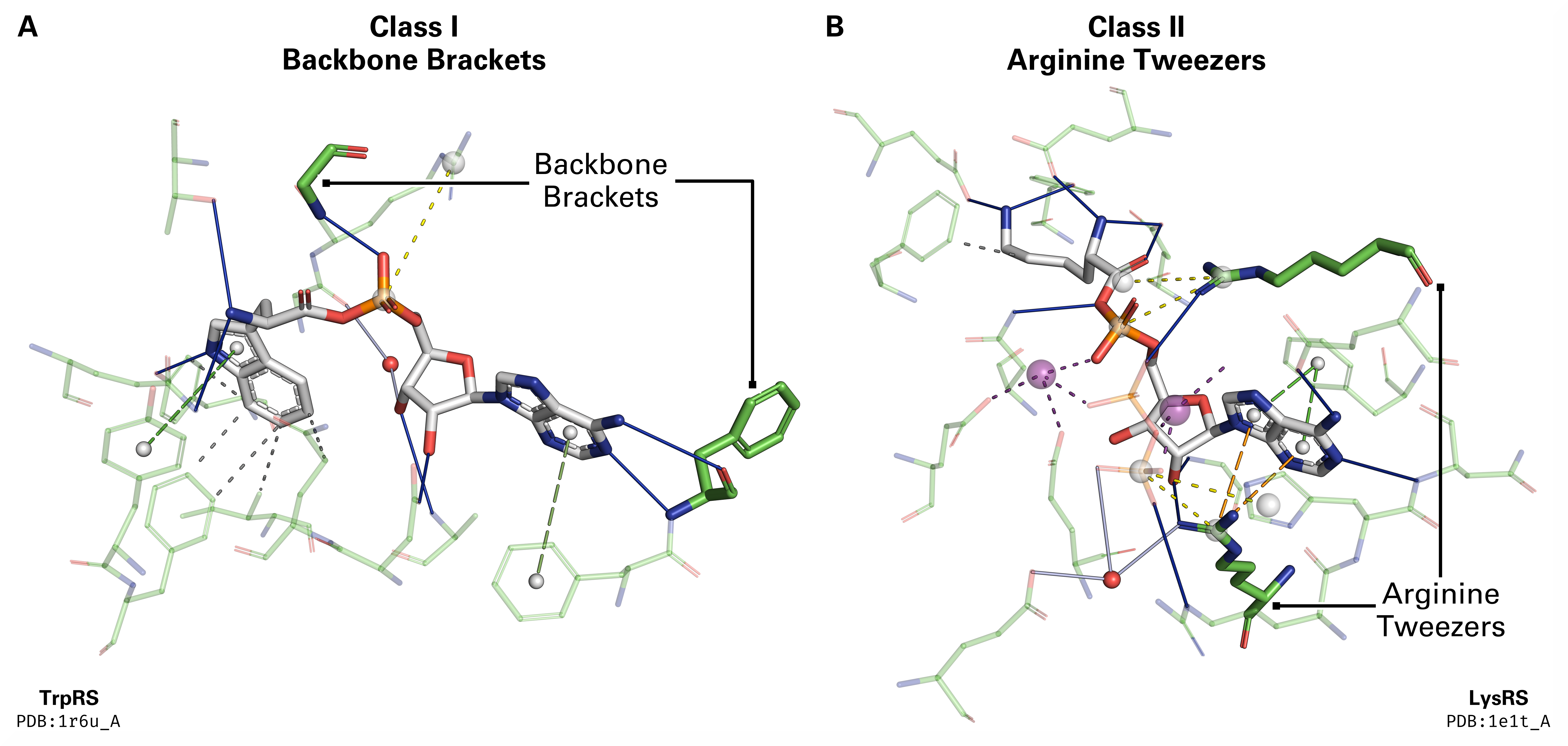

### Supplementary Materials

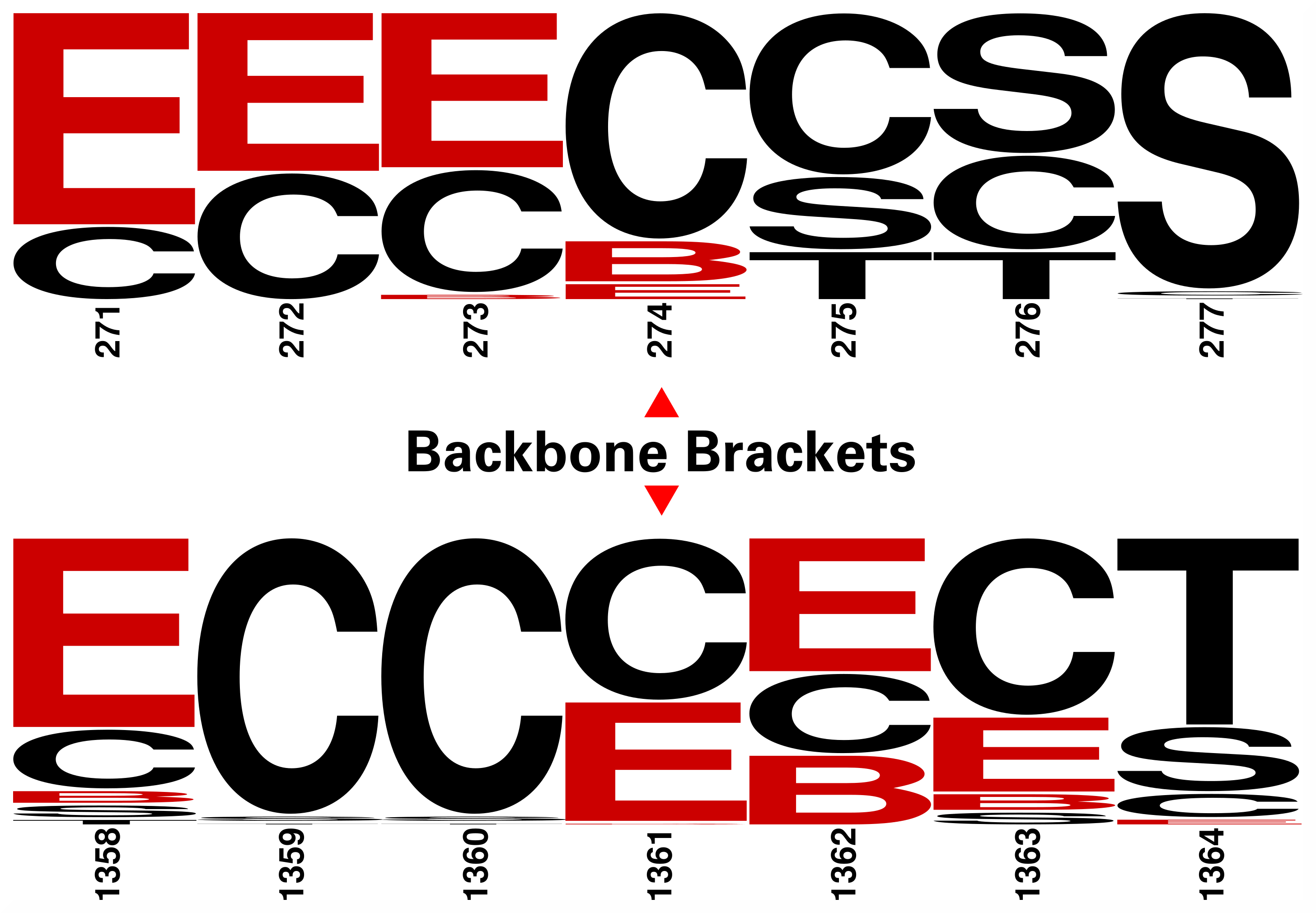

### Supplementary Materials

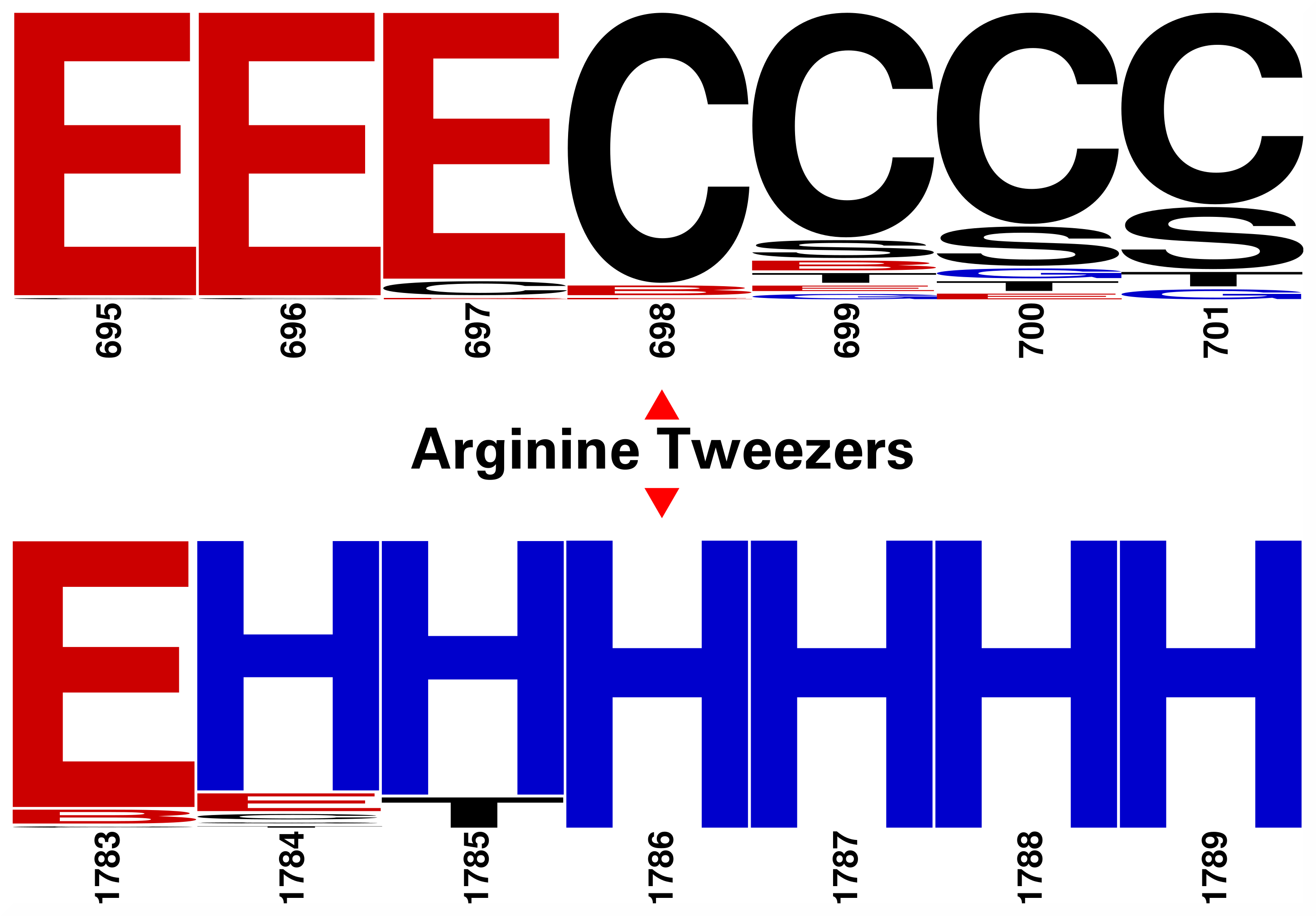

### Supplementary Materials

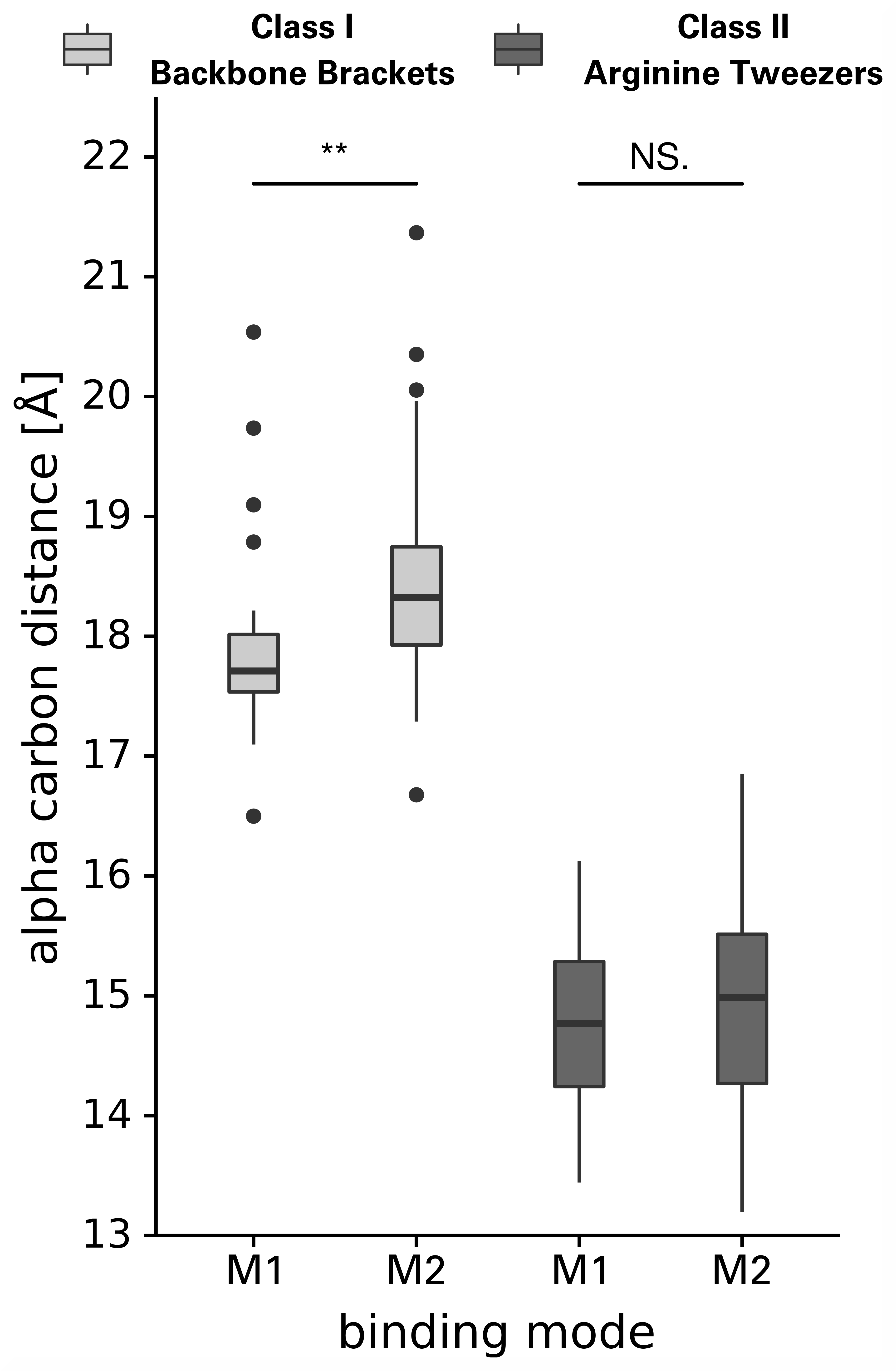
